# Early Corneal Host-Response Signatures Associate with Disease Severity and Etiology-Related Clinical Outcomes in Fungal Keratitis

**DOI:** 10.64898/2026.11.23.768774

**Authors:** Swagata Ghosh, Anusha Erulandi, Dhanu Prakash Muthupandi, Abinaya Krishnan, Mummaneni Sahana, Ramalakshmi Raman, Prajna Lalitha, N. Venkatesh Prajna

## Abstract

**Background:** To advance host-based predictive marker development for individual risk assessment in fungal keratitis (FK), we investigated corneal host-response gene-expression signatures associated with etiology-related disease course and clinical outcomes.

**Methods:** We compared severity, healing, and visual outcomes in 1,304 patients with *Fusarium* spp., *Aspergillus flavus*, or culture-negative keratitis. Presentation-stage corneal scrapings from 70 prospective patients underwent bulk RNA sequencing stratified by etiology, severity, and outcome. Candidate signatures were independently validated by RT-qPCR in 102 patients using clinical correlations, individual cutoff-based profiling, and masked testing, followed by experimental evaluation in corneal epithelial and macrophage-like cells challenged with live fungal spores.

**Results:** *Fusarium* keratitis more frequently healed with preserved vision. *A. flavus* and culture-negative keratitis had the highest and lowest proportions of severe and unhealed disease, respectively, despite similarly poor visual outcomes among healed patients. Transcriptomics paralleled these patterns, revealing a distinct *Fusarium* host response but substantial *A. flavus*-culture-negative concordance. A coordinated 8-gene innate-response module correlated with ulcer size and discriminated severe from non-severe disease (AUC, 0.82–0.96), while showing more robust expression in *A. flavus* and culture-negative than *Fusarium* keratitis. Higher corneal *ICOSL* was associated with healing, whereas increased *SLC26A4*/pendrin characterized non-healing and visual impairment, particularly in *Fusarium* keratitis. Together, these formed a 10-gene panel whose cutoff-based profiles showed concordance with severity and outcome in masked testing. Experimental evaluation revealed cell- and pathogen-specific expression patterns consistent with patient profiles.

**Conclusions:** Presentation-stage corneal host-response profiles capture etiology-associated differences in FK severity and outcome and identify a candidate 10-gene framework for individualized risk stratification.

**Graphical Abstract:** 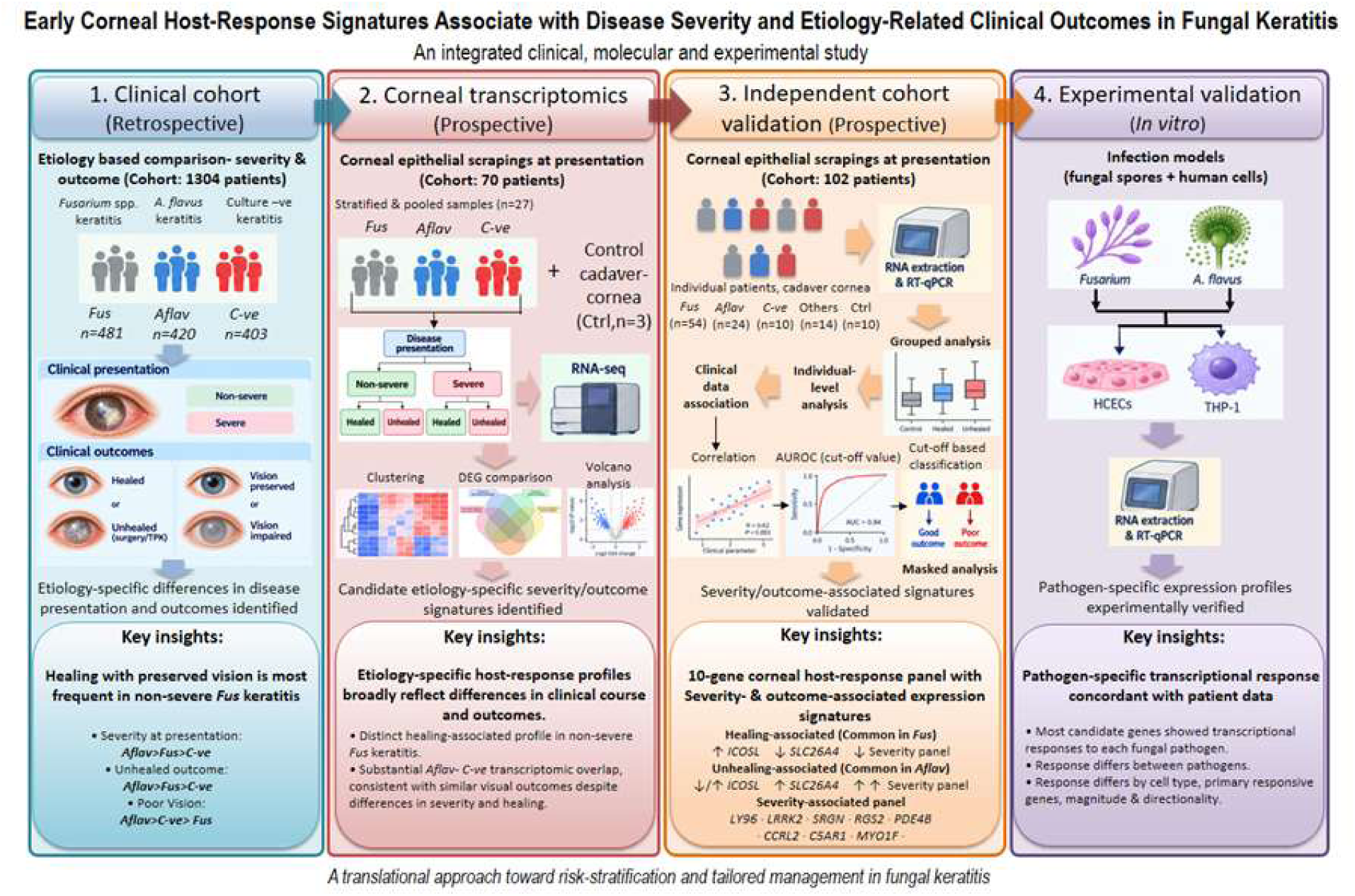

## Introduction

Blindness due to corneal opacity by microbial keratitis (MK) disproportionately affects working-age populations in low- and middle-income tropical countries ^1,2^. Fungal keratitis (FK), accounting for nearly half of MK cases, is associated with worse clinical outcomes caused primarily by filamentous fungi like *Fusarium* and *Aspergillus* ^3–5^. Despite standard antifungal therapy, outcomes remain poor, with approximately one-quarter of cases requiring therapeutic penetrating keratoplasty or resulting in severe visual impairment ^1,5,6^. Moreover, the lack of objective tools for early risk stratification and growing antifungal resistance highlight the need for improved prognostic and host-directed therapeutic strategies in FK ^4,7^.

Host-mediated inflammation is increasingly recognized as a major determinant of tissue injury in FK. Human corneal studies demonstrate neutrophil-dominated innate responses and matrix metalloproteinase activation in advanced disease, supporting inflammation-driven stromal damage ^8,9^, while miRNA profiling in *Fusarium* keratitis implicates dysregulated TLR, NF-κB, IL-17, and TNF signaling in disease progression and severity ^10^. Importantly, ocular-surface transcriptomics and tear proteomics demonstrate host-response activation at disease onset ^11–13^. Epidemiological studies further suggest variation in clinical outcomes across fungal etiologies ^14^, potentially reflecting both shared and distinct host-response pathways ^8,15^. However, these early responses remain insufficiently defined at the gene-expression level or linked to etiology, subsequent disease course, and treatment outcomes. Comparing presentation-stage corneal profiles across etiologies and severity- and outcome-stratified patients could therefore identify beneficial and detrimental host-response signatures with potential for early risk assessment.

Here, we first characterized disease course, treatment and visual outcomes across a large retrospective cohort of patients with *Fusarium* spp., *Aspergillus flavus*, and culture-negative keratitis. We then investigated whether these clinically defined phenotypes were reflected in distinct corneal host-response programs by prospectively profiling gene expression in corneal scrapings collected at presentation from severity- and outcome-stratified patients. Independent patient cohorts and experimental models were used to validate clinically relevant gene signatures and assess their association with disease severity and treatment and visual outcomes. This integrated clinical and transcriptomic approach identified early corneal host-response signatures with potential utility for prognostic assessment and risk stratification in FK.

## Methods

### Retrospective study Design

Participants were selected retrospectively from the patients treated during the period of year 2021-2024 at the Cornea Clinic, Aravind Eye Hospital, Madurai. Cohort construction is detailed in Supplementary Methods.

### Prospective study participants and Sample Collection

The study adhered to the tenets of the Declaration of Helsinki and was approved by the Institutional Ethics Committee of Aravind Eye Hospital (Project code: IRB2021016BAS). Written informed consent was obtained from all participants. Study participants were recruited at first presentation at the Cornea Clinic, Aravind Eye Hospital, Madurai between the period 2021 - 2026. Corneal tissue surrounding the ulcer was collected prior to treatment with a sterile corneal swab and scraper and placed into tubes containing Zymo DNA/RNA Shield buffer (Zymo Research, Irvine, CA) and stored at −80°C until total RNA isolation.

### Cell culture, LPS stimulation and infection with *Fusarium* spp. or *A. flavus* spore

Recombinant SV-40 adenovirus vector immortalized human corneal epithelial cell line RCB2280 (HCEC) was purchased from Riken cell bank, Japan and Tohoku Hospital pediatrics-1 (THP-1) cell lines were obtained from the National Center for Cell Sciences (NCCS), Pune are used for the in vitro experiment detailed in Supplementary Methods.

#### RNA extraction

Total RNA was extracted from patient samples and cultured cells using TRIzol-Chloroform method detailed in Supplementary Methods.

### RNA sequencing

Patient derived total RNA from different disease groups and controls were outsourced to Biokart India Pvt. Ltd. for RNA sequencing on an illumina platform detailed in Supplementary Methods.

### RT-qPCR Analysis

For all samples, total RNA was used for cDNA synthesis (QuantiTect Reverse Transcription Kit; Qiagen, Hilden, Germany). Quantitative real-time PCR was performed using the Quantinova SYBR Green PCR Kit (Qiagen) in Quantstudio Real-Time PCR system (Applied Biosystems, Waltham, MA) using previously reported oligonucleotides (Supplementary Table 1).RT-qPCR analysis is detailed in Supplementary Methods.

### Statistical Analysis

All statistical methods used in retrospective and prospective studies are detailed in Supplementary Methods.

## Results

### Clinical presentation and outcomes differ by fungal etiology

We first examined etiologic differences in presentation severity and clinical outcomes among 1,304 patients with *Fusarium* spp. (n=481), *Aspergillus flavus* (n=420), or culture-negative keratitis (n=403). Sex distribution and history of ocular trauma were similar across groups, whereas age and systemic comorbidities differed (Supplementary Table 2).

Disease severity at presentation differed by etiology (*P*<0.001), with severe disease more frequent in *A. flavus* (49.8%) than *Fusarium* (26.4%) or culture-negative keratitis (23.3%) (Supplementary Table 3, Supplementary Figure 1B). Because the cohorts constructed for etiologic comparisons did not reflect the relative frequency of these infections in clinical practice, we examined a separate 3-month consecutive cohort that preserved the natural distribution of etiologies among presenting cases. Although the overall difference in presentation severity across etiologies did not reach significance (*P*=0.077), *A. flavus* remained more likely to present with severe disease than both *Fusarium* (*P*=0.035) and culture-negative keratitis (*P*=0.025), whereas *Fusarium* and culture-negative keratitis did not differ (*P*=0.956) (Supplementary Table 4).

Treatment outcomes showed a clearer etiologic pattern. An unhealed outcome requiring therapeutic penetrating keratoplasty occurred in 40.5% of *A. flavus*, 22.0% of *Fusarium*, and 9.4% of culture-negative cases (*P*<0.001; Supplementary Table 3, Supplementary Figure 1C), and the higher frequency of treatment failure with *A. flavus* was reproduced in the consecutive cohort (*P*<0.001; Supplementary Table 4). Because *A. flavus* also presented with greater severity in the retrospective cohort, outcomes were examined within severity strata. Among non-severe cases, *A. flavus* had more unhealed outcomes than *Fusarium* (*P*=0.005) and culture-negative keratitis (*P*<0.001) (Table 1, Supplementary Figure 1D). Among severe cases, outcomes did not differ significantly between *A. flavus* and *Fusarium*, whereas treatment failure remained more frequent in these groups compared to culture-negative keratitis (Table 1, Supplementary Figure 1E).

**Table 1.** Comparison of clinical and visual outcomes across etiologies of keratitis.

| **Keratitis** | **Presentation<sup>a</sup>**<br>**(*P*<0.001<sup>g</sup>)** | **Outcome<sup>b</sup>**<br>**(*P*<0.001<sup>g</sup>)** |  | **OR (95% CI)<sup>e</sup>** | **P Value** | **OR (95% CI)<sup>e</sup>** | **P Value** | **Healed- Vision<sup>c</sup>**<br>**(*P*<0.001<sup>g</sup>)** |  | **OR (95% CI)<sup>f</sup>** | **P Value** | **OR (95% CI)<sup>f</sup>** | **P Value** |
|---|---|---|---|---|---|---|---|---|---|---|---|---|---|
| *A. flavus*<br>(n=420) | Non-severe<br>(50.2%) | Healed<br>(80.1%) | Unhealed<br>(19.9%) | 1.95<br>(1.22- 3.13) | 0.005 | 7.43<br>(3.64-15.19) | <0.001 | GV<br>(68.1%) | PV<br>(31.9%) | 2.67<br>(1.70-4.17) | <0.001 | 1.29<br>(0.85-1.94) | 0.233 |
|  | Severe<br>(49.8%) | Healed<br>(38.8%) | Unhealed<br>(61.2%) | 1.46<br>(0.94-2.28) | 0.096 | 3.72<br>(2.21-6.28) | <0.001 | GV<br>(24.7%) | PV<br>(75.3%) | 4.11<br>(2.01-8.40) | 0.001 | 1.63<br>(0.80-3.34) | 0.18 |
| *Fusarium sp.*<br>(n=481) | Non-severe<br>(73.6%) | Healed<br>(88.7%) | Unhealed<br>(11.3%) | Ref.<sup>d</sup> | - | 3.81<br>(1.87-7.75) | <0.001 | GV<br>(85.0%) | PV<br>(15.0%) | Ref.<sup>d</sup> | - | 0.48<br>(0.32-0.72) | <0.001 |
|  | Severe<br>(26.4%) | Healed<br>(48%) | Unhealed<br>(52%) | Ref.<sup>d</sup> | - | 2.55<br>(1.45-4.48) | 0.001 | GV<br>(57.4%) | PV<br>(42.6%) | Ref.<sup>d</sup> | - | 0.40<br>(0.19-0.81) | 0.012 |
| Culture -ve<br>(n=403) | Non-severe<br>(76.7%) | Healed<br>(96.8%) | Unhealed<br>(3.24%) | 0.26<br>(0.13-0.53) | <0.001 | Ref.<sup>d</sup> | - | GV<br>(73.2%) | PV<br>(26.8%) | 2.08<br>(1.39-3.10) | <0.001 | Ref.<sup>d</sup> | - |
|  | Severe<br>(23.3%) | Healed<br>(70.2%) | Unhealed<br>(29.8%) | 0.39<br>(0.22- 0.69) | 0.001 | Ref.<sup>d</sup> | - | GV<br>(34.9%) | PV<br>(65.1%) | 2.52<br>(1.23-5.15) | 0.012 | Ref.<sup>d</sup> | - |

Among healed patients, however, the etiologic pattern for visual outcome differed from that for clinical healing. Overall, *Fusarium* keratitis had the greatest proportion of patients with good visual outcome compared with both *A. flavus* and culture-negative keratitis (Supplementary Table 5, Supplementary Figure 1F), a pattern that persisted across severity strata (Table 1, Supplementary Figure 1G & 1H). Compared with *Fusarium*, poor vision was more frequent with *A. flavus* in both non-severe (*P*<0.001) and severe disease (*P*=0.001) (Table 1). Notably, despite its higher healing rate, culture-negative keratitis had visual outcomes comparable to *A. flavus* across severity strata (Table 1, Supplementary Figure 1G & 1H). Because central ulcer location could independently contribute to poor vision, we assessed ulcer position in a random subset of 30 patients; central involvement was not greater in *A. flavus*, suggesting that ulcer location did not account for the observed visual differences (Supplementary Table 6).

### Corneal transcriptomic profiles reflect etiology-associated severity and outcome patterns

To examine if corneal transcriptional programs at disease presentation underlie the etiology based differences in disease severity and outcomes, we performed RNA sequencing in prospectively sampled patient-derived corneal scrapings. Patient samples were stratified by etiology, clinical severity (non-severe, severe) and outcome (healed, unhealed) with cadaver cornea serving as healthy controls. While all presentation-outcome categories were represented in *Fusarium* keratitis, partial categories could be assembled for *Aspergillus flavus* and culture-negative cases due to natural differences in occurrence of specific clinical groups in these etiologies (Figure 1A). Transcriptomic profiles of the available clinical groups separated them into clusters strikingly consistent with their clinical stratifications as demonstrated by Principal Component analyses (Figure 1B and Supplementary Figure 2A). Within infected samples, non-severe healed cases formed 2 distinct clusters, with *Fusarium* keratitis forming one unique group separated from *A. flavus* and culture-negative profiles overlapping into another cluster (Figure 1B). The severe-healed and unhealed profiles were separate from these two clusters, overlapped considerably while indicating some degree of segregation between *Fusarium* and the other two etiologies within this cluster (Figure 1B).

**Figure 1.**
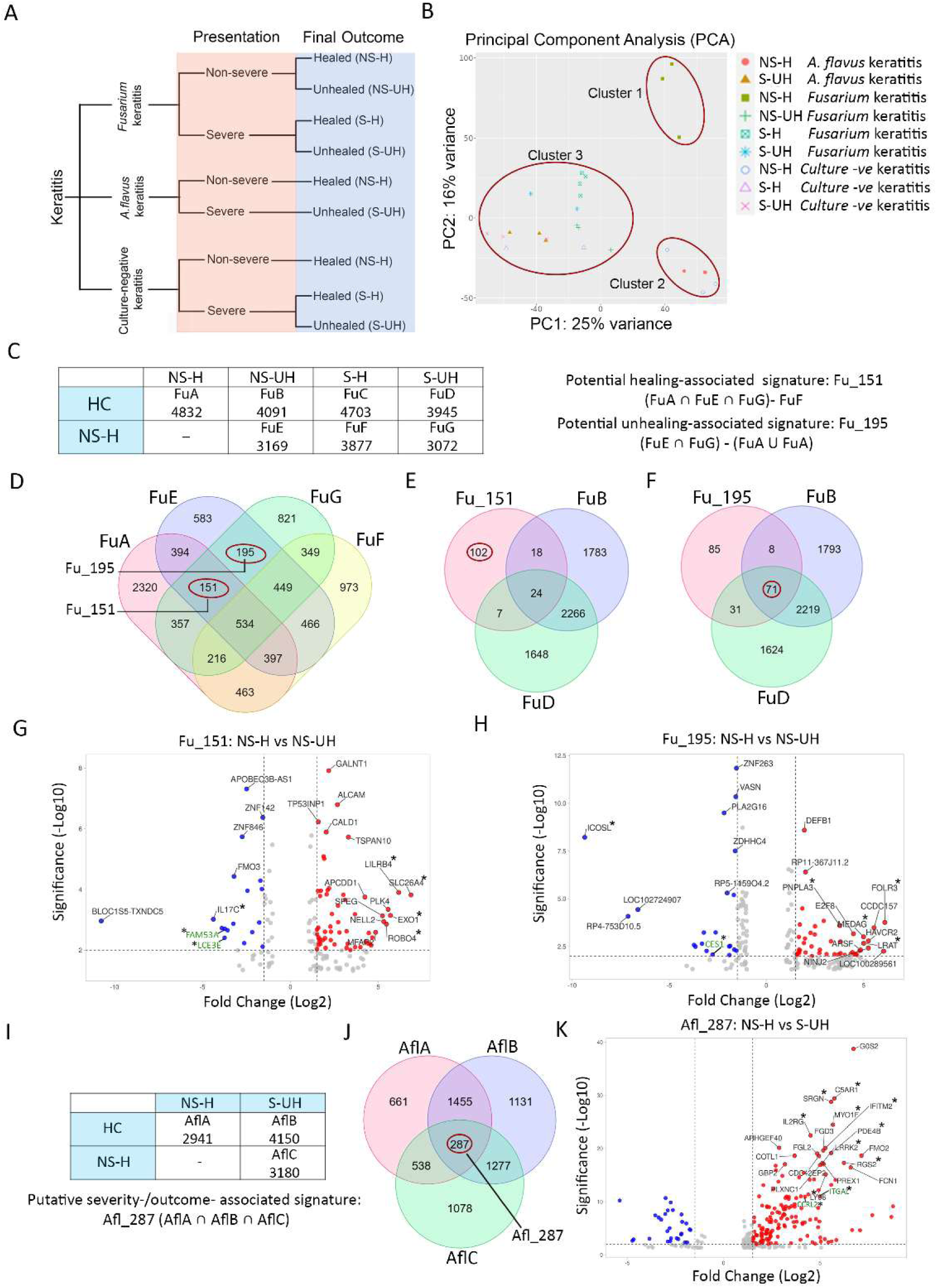
Transcriptomic profiling reveals severity- and outcome-associated gene signatures in fungal keratitis. RNA-seq grouping of fungal keratitis samples by etiology, presentation severity, and final clinical outcome (A). PCA showing clustering of transcriptomic profiles across clinical groups (B). Differential-expression comparison strategy used to derive the potential *Fusarium* healing-associated (Fu_151) and unhealing-associated (Fu_195) signatures (C). Venn analysis defining Fu_151 and Fu_195 (D), followed by comparison with *Fusarium* control-versus-unhealed DEG sets to identify 102 healing-specific genes and 24 shared DEGs within Fu_151 (E), and a common 71-gene unhealing-associated signature within Fu_195 (F). Volcano plots of Fu_151 (G) and Fu_195 (H) genes comparing non-severe healed (NS-H) with non-severe unhealed (NS-UH) disease; top 20 DEGs are labeled in black, additional candidates in green, and prioritized validation candidates by asterisks. Differential-expression comparison strategy used to derive a putative *A. flavus* severity-/outcome-associated signature (I), with Venn analysis identifying the 287-gene Afl_287 set (J). Volcano plot of Afl_287 genes comparing NS-H with severe-unhealed (S-UH) disease; top 20 DEGs are labeled in black, additional candidates in green, and prioritized validation candidates by asterisks (K). HC, healthy control; DEG, differentially expressed gene.

Analysis across presentation-outcome subtypes (Supplementary Methods) within *Fusarium* keratitis allowed us to identify outcome associated differentially expressed genes (DEGs), irrespective of severity status at presentation. Importantly, DEG analysis in *Fusarium* keratitis pointed to distinct subsets of genes associated with healing and unhealing outcomes: 151 genes comprising a healing-associated signature and 195 genes comprising an unhealing-associated signature (Figure 1C and 1D). These signatures were supported by hierarchical clustering analysis of normalized gene expression levels using an unsupervised approach (Supplementary Figure 3 and 4). Healing associated signatures were those significant gene-expression changes that occurred primarily in the non-severe healed groups relative to the controls and did not occur in the unhealed groups and hence were significantly altered in the unhealed groups relative to the healed (Figure 1C-1E, Supplementary Figure 3). Similarly unhealing associated signatures were significantly and uniquely altered in the unhealed groups and not the healed groups relative to the controls and hence differed significantly between non-severe healed and the unhealed groups (Figure 1C, 1D, 1F and supplementary Figure 4). We observed minimal overlap between common DEGs in disease groups from healthy control based analysis and from analysis relative to the non-severe healed group (Supplementary Figure 2B-2D).

Candidate genes were selected from the 151 DEGs based on fold change and level of significance in non-severe healed group relative to both non-severe and severe unhealed groups and healthy controls (Figure 1G, Supplementary Figure 2E, Supplementary Table 7). These include *IL17C, FAM53A, LCE3E, SLC26A4, EXO1, ROBO4, LILRB4* (Figure 1G, Supplementary Figure 2E and Supplementary Table 7). While all of these gene expression changes were specific to healing (102 genes, Figure 1E), *LILRB4*, was the only one gene to be downregulated in healed and upregulated in unhealed samples relative to controls. Similar to the healing signature candidates, unhealing signature candidates were identified within the 195 DEGs based on fold change and level of significance in both non-severe and severe unhealed groups relative to the non-severe healed group and controls (Figure 1H, Supplementary Figure 2F, Supplementary Table 8). These included *CES1, ICOSL, FOLR3, LRAT, MEDAG* and *PNPLA3* (Figure 1H, 1F, Supplementary Figure 2F and Supplementary Table 8) representing the unhealing-specific gene expression changes relative to the controls (71 genes, Figure 1F).

In contrast with *Fusarium* keratitis analysis, *A. flavus* keratitis transcriptome allowed identification of a subset of DEGs associated with either severity status at presentation or treatment outcome, or both, since only non-severe healed and severe unhealed clinical groups were available. As opposed to *Fusarium* keratitis, where healing and unhealing associated gene subsets were distinct, we identified a common subset of 287 genes differentially expressed in both the clinical groups as compared with the control and relative to each other (Figure 1I and 1J). Unsupervised hierarchical clustering analysis further supported that clinical categories of *A. flavus* keratitis can be clustered by distinct expression profiles of these 287 genes (Supplementary Figure 5). In addition, non-severe healed group of the culture negative keratitis shared these 287 DEGs with similar profile as the non-severe healed *A. flavus* keratitis as revealed by clustering analysis (Supplementary Figure 5). Once again candidate genes were selected based on fold change and significance in all three comparisons within *A. flavus* keratitis and these include *C5AR1, RGS2, CCRL2, IL2RG, SRGN, LY96, LRRK2, IFITM2, SLC16A3, PDE4B, MYO1F,* and *ITGAL* (Figure 1K and Supplementary Table 9).

Comparative analysis of three etiologies point to the significant overlap of gene-expression profile between *A. flavus* keratitis and the culture negative group (Supplementary Figure 6). DEG comparison within culture negative keratitis showed that common DEGs identified from control based analysis has limited overlap with common DEGs identified from non-severe healed based analysis. In spite of this, the overlapping subset included several of those DEGs identified as severity-outcome associated genes in *A. flavus* keratitis transcriptome and were identified to be top candidates in culture negative keratitis transcriptome as well (Supplementary Figure 6A-6D). In fact, non-severe healed based DEGs from culture negative keratitis and *A. flavus* keratitis showed a large overlap (Supplementary Figure 6E), such that there were 65% of the top predicted genes differentially expressed in severe-unhealed group matched between the two etiologies in the relative to the non-severe healed group (Supplementary Figure 6F and 6G). Healing and unhealing signatures identified in *Fusarium* keratitis analysis were not found among the most significant alterations in the other two etiology groups. Instead, several of the severity-outcome associated genes for *A. flavus* and culture negative keratitis belong to a subset of genes that differentiate the *Fusarium* keratitis non-severe healed group from that of the other two etiologies (Supplementary Figure 6H and 6I), pointing at the distinctiveness of the *Fusarium* keratitis healing-associated corneal transcriptional program.

### Independent validation identifies severity- and outcome-associated host-response profiles

Candidate genes identified by RNA sequencing were evaluated by RT-qPCR in an independent patient cohort using individual corneal scrapings collected at presentation, with healthy donor corneal scrapings as controls (Supplementary Table 10). Of 15 *Fusarium*-derived candidates evaluated, 8 were detectable, of which *ICOSL, SLC26A4, IL17C, CREB5,* and *PDE4B* were differentially expressed relative to controls (Figure 2A–E). *ICOSL* expression was higher in healed than unhealed *Fusarium* keratitis, whereas *SLC26A4* was increased in unhealed disease. *IL17C, CREB5*, and *PDE4B* were elevated in both outcome groups, with *PDE4B* showing a significant further increase in unhealed cases. *LILRB4, PNPLA3,* and *FAM53A* were not significantly different from controls (Supplementary Figure 7A-7C). Notably, higher *ICOSL* expression above a threshold, derived by a healed-versus-unhealed AUROC analysis (Figure 3A), associated significantly with healed outcome (Figure 3B) while it did not correlate with presentation severity and ulcer size (Figure 3C).

**Figure 2.**
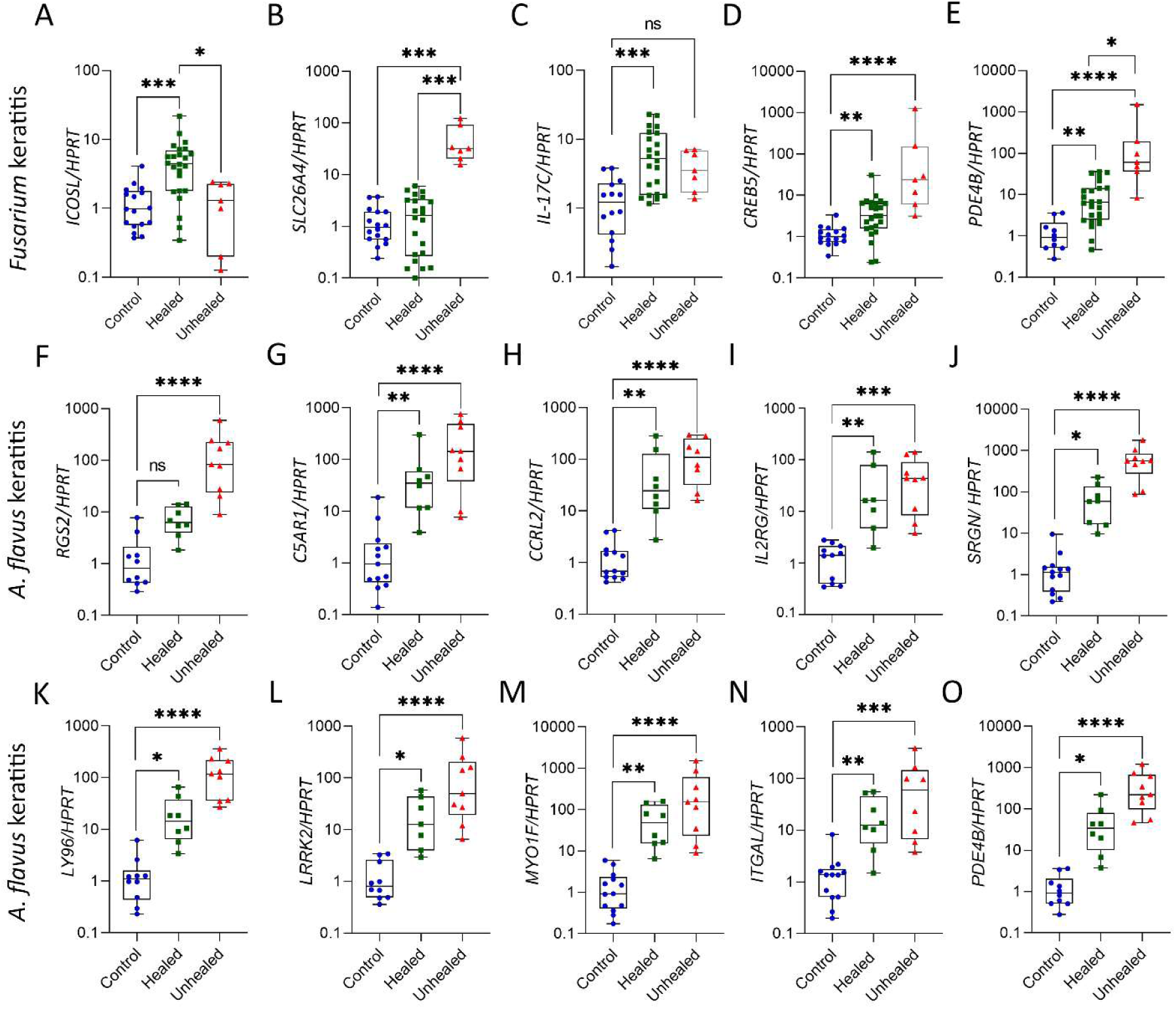
Validation of gene expression profiles in patients with *Fusarium* and *Aspergillus flavus* keratitis. HPRT-normalized expression of candidate genes in corneal samples from patients with *Fusarium* (A-E) and *A. flavus* (F-O) keratitis, shown as fold change relative to healthy cadaveric controls. Box plots show the median and interquartile range with individual data points for control, healed, and unhealed groups. Statistical significance was determined by Kruskal-Wallis test with Dunn’s multiple-comparisons test: \**P*<.05; \*\**P*<.01; \*\*\**P*<.001; \*\*\*\**P*<.0001; ns, nonsignificant.

**Figure 3.**
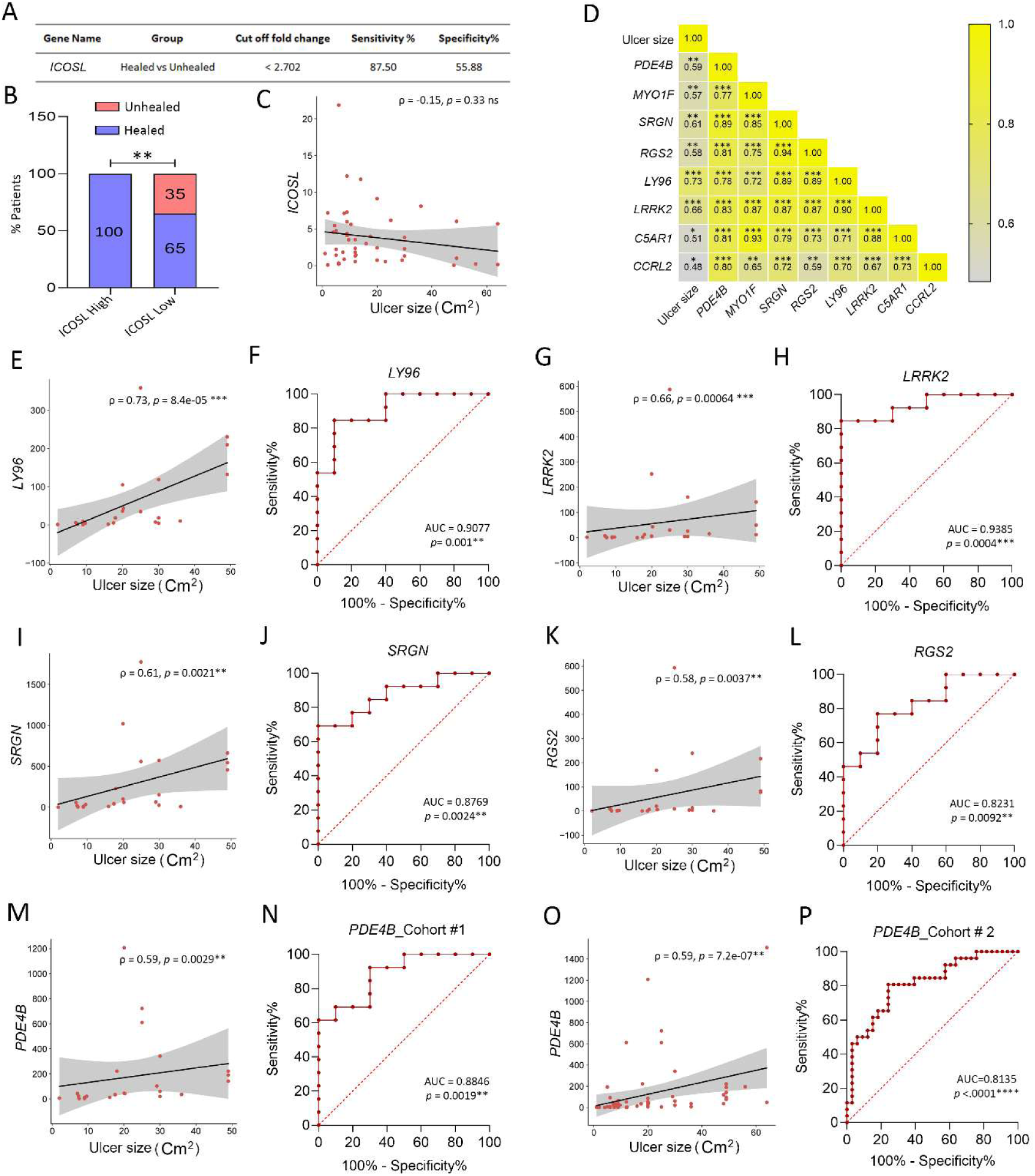
Association of presentation-stage host-response gene expression with clinical parameters. In *Fusarium* keratitis (n=37), ROC analysis distinguishing unhealed from healed disease identified *ICOSL* <2.702-fold as the optimal cutoff for unhealed disease (87.5% sensitivity; 55.88% specificity) (A). All patients with high *ICOSL* expression (>2.702; 17/17) healed with medical therapy, compared with 65% (13/20) with low expression (<2.702; Fisher exact test, *P*<.01) (B); *ICOSL* expression did not correlate with ulcer size (C). Spearman correlation matrix of ulcer size and the 8-gene severity-associated panel (n=23; *Fusarium*, n=9; *A. flavus*, n=14) (D). Correlations of ulcer size with *LY96* (E), *LRRK2* (G), *SRGN* (I), *RGS2* (K), and *PDE4B* (M), with corresponding ROC analyses distinguishing severe from non-severe disease (F, H, J, L, N). *PDE4B* correlation with ulcer size (O) and ROC analysis for severe versus non-severe disease (P) were further evaluated in the larger validation cohort (n=59; *Fusarium*, n=38; *A. flavus*, n=21). Spearman correlation coefficients (ρ), AUCs, and *P* values are shown; \**P*<.05, \*\**P*<.01, \*\*\**P*<.001, \*\*\*\**P*<.0001.

Validation of the *A. flavus* signature revealed broader upregulation of the prioritized host-response genes. *C5AR1, CCRL2, IL2RG, SRGN, LY96, LRRK2, MYO1F, ITGAL,* and *PDE4B* were significantly increased relative to controls, with most elevated in both healed and unhealed disease (Figure 2F–2O). *RGS2* was significantly increased only in unhealed cases, whereas *IFITM2* and *SLC16A3* were unchanged (Supplementary Figure 7D & 7E). Although grouped analysis did not significantly separate healed from unhealed *A. flavus* keratitis, several genes showed increasing expression with greater disease severity, prompting us to examine their relationship with clinical parameters at the individual-patient level.

Across *A. flavus* and *Fusarium* keratitis, *PDE4B, MYO1F, SRGN, RGS2, LY96, LRRK2, C5AR1,* and *CCRL2* showed significantly correlated expression, correlated with ulcer size, and discriminated severe from non-severe disease (AUC, 0.82–0.96; Figure 3D–3P; Supplementary Table 11), defining a coordinated severity-associated expression profile. We then applied the AUROC-derived severity thresholds to individual expression profiles and examined their relationship with subsequent outcomes. High expression across multiple severity-associated genes predominated among patients with poor clinical and visual outcomes, particularly in *A. flavus*, whereas healed *Fusarium* cases generally showed limited expression of this program (Supplementary Tables 12 and 13). Similar patient-level patterns were observed in other etiologies: favourable clinical and visual outcomes in *Curvularia* and *Pseudomonas aeruginosa* keratitis were generally accompanied by limited severity-associated gene expression, whereas culture-negative keratitis more closely resembled the higher-expression profiles observed in *A. flavus* (Supplementary Table 14; Supplementary Figures 8 and 9). Importantly, Higher *ICOSL* with limited severity-associated expression more frequently associated with favourable clinical and visual outcomes across different etiologies, primarily in *Fusarium* keratitis (Figure 3A & 3B; Supplementary Table 13 and 14). Low *ICOSL* levels combined with low to moderate levels of the severity-associated genes were associated with variable outcomes across different etiologies, identifying patients not resolved by the current signatures (Supplementary Table 13 and 14).

Finally, we explored these expression patterns in six samples analyzed while masked to pathogen identity and clinical characteristics. We found that expression-based severity assignment was concordant with presentation severity in five of six cases, while outcome profiles reproduced the favourable, unfavourable, and indeterminate patterns observed in the validation cohort (Table 2; Supplementary table 15 and 16). Together, these analyses link presentation-stage host gene expression with disease severity and subsequent clinical course, while identifying candidate patient-level profiles for prospective validation and outcome-based risk stratification.

**Table 2.** Masked evaluation of presentation-stage host-response profiles and concordance with clinical course. . Expression profiles were interpreted while masked to pathogen identity and clinical characteristics using the candidate framework in Supplementary Table 16.

| Patient | 10-gene host-response profile at presentation*† | Masked severity interpretatio‡ | Masked outcome interpretation‡ | Actual severity | Clinical course at last follow-up | Pathogen | Clinical Concordance |
|---|---|---|---|---|---|---|---|
| **U1** | ICOSL − \| SLC26A4 + \| 1/8 ↑ | NS | Fus: UH; Aflav: UFC, Var; C–:H | S | TPK | Fus | Severity ✕ / Outcome ✓ |
| **U2** | ICOSL − \| SLC26A4 − \| 6/8 ↑ | S | Fus/Aflav: UH; C–: UFC, H | S | TPK | Aflav | Severity ✓/ Outcome ✓ |
| **U3** | ICOSL + \| SLC26A4 + \| 1/8 ↑ | NS | Fus/Aflav/C−: H | NS | Healed, good vision | C– | Severity ✓/ Outcome ✓ |
| **U4** | ICOSL + \| SLC26A4 + \| 2/8↑, 6/8 ↑↑ | S | Fus: UFC, Var; Aflav: UH; C−: H | S | Persistent, Non-healing (LOF) | Fus | Severity ✓/ Disease course ✓ |
| **U5** | ICOSL + \| SLC26A4 + \| 3/8↑, 5/8 ↑↑ | S | Fus: UFC, Var; Aflav: UH; C−: H | S | Healed, scar with vision | C– | Severity ✓/ Outcome ✓ |
| **U6** | ICOSL − \| SLC26A4 + \| 6/8 ↑ | S | Fus: UH; Aflav: UH; C−: UFC, H | S | Healed, scar, good vision | Other /mixed | Severity ✓ Outcome-NA |

**Abbreviations:** Aflav, *Aspergillus flavus*; C−, culture-negative; Fus, *Fusarium* spp.; H, healed; LOF, loss to follow-up; NA, not applicable; NS, non-severe; S, severe; TPK, therapeutic penetrating keratoplasty; UFC, unfavourable course; UH, unhealed; Var, variable/indefinite.
**Footnotes:**
\* + and − indicate expression above and below the predefined *ICOSL* and *SLC26A4* cutoffs, respectively.
† For the 8-gene severity-associated panel, ↑ indicates expression above the gene-specific cutoff and ↑↑ indicates >5-fold above cutoff; *n*/8 indicates the number of genes meeting the corresponding threshold.
‡ Masked severity was interpreted from the number and magnitude of elevated severity-associated genes. Masked outcome interpretations were assigned for each candidate etiology and subsequently compared with the revealed pathogen and clinical course.

### Experimental Models Reveal Pathogen-Specific Responses Concordant With Patient Profiles

To determine whether the host-response signatures identified in patients were directly responsive to inflammatory and fungal challenge, we examined validated candidates in human corneal epithelial cells (HCECs) and differentiated THP-1 macrophage-like cells. Responses were first assessed following LPS stimulation (Supplementary Figures 10 and 11) and then following infection with clinical isolates of *Fusarium* spp. or *A. flavus* (Figures 4 and 5; Supplementary Figure 12).

**Figure 4.**
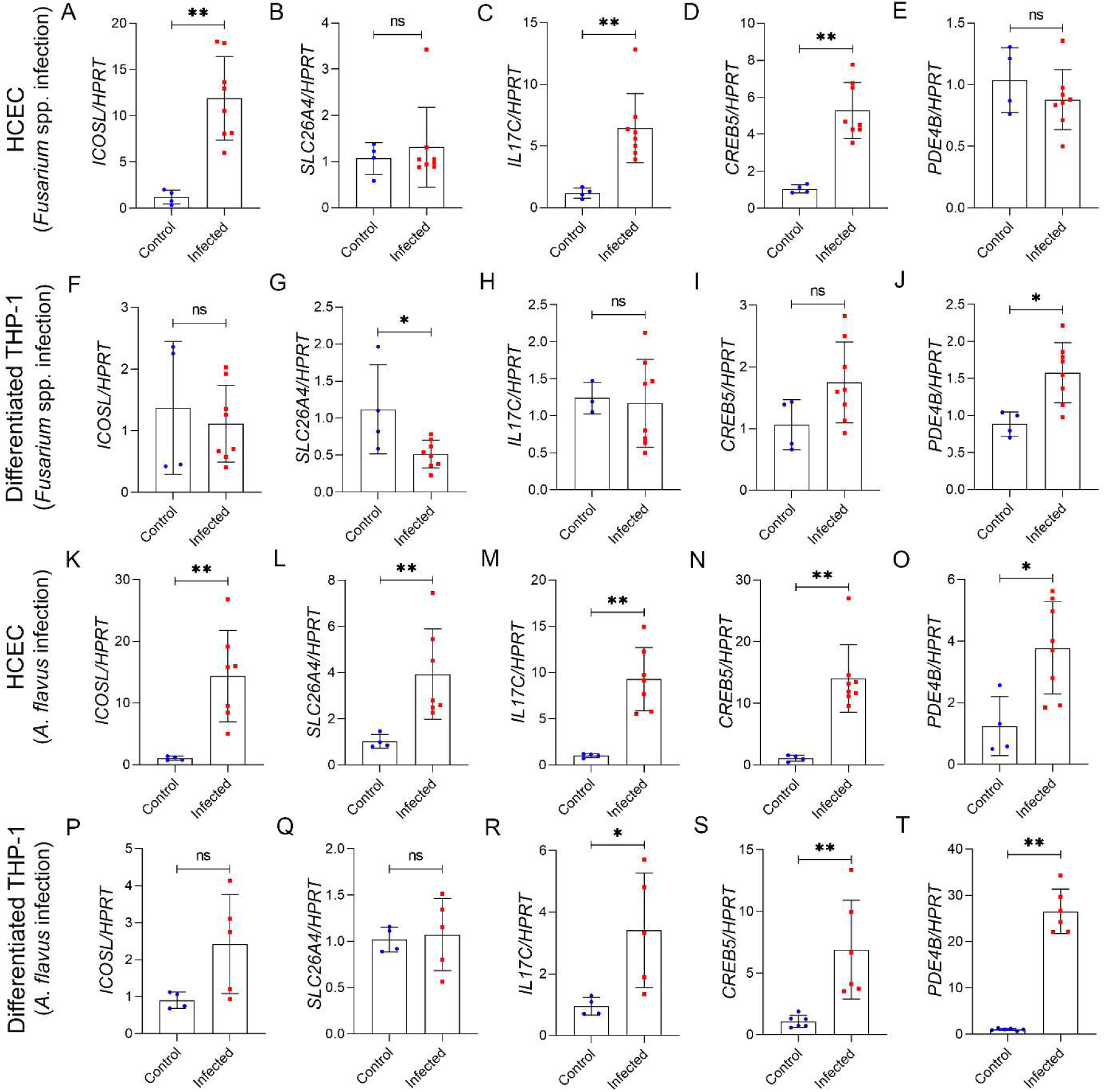
Expression of *Fusarium* keratitis-associated genes in HCECs and differentiated THP-1 cells following fungal spore infection. HPRT-normalized expression of candidate genes in human corneal epithelial cells (HCECs) and differentiated THP-1 cells infected with *Fusarium* spores (A-J) or *A. flavus* spores (K-T), shown as fold change relative to uninfected controls. Bars represent mean ± SD with individual data points representing biological replicates. Statistical significance was determined by Mann-Whitney U test: \**P*<.05; \*\**P*<.01; ns, nonsignificant.

**Figure 5.**
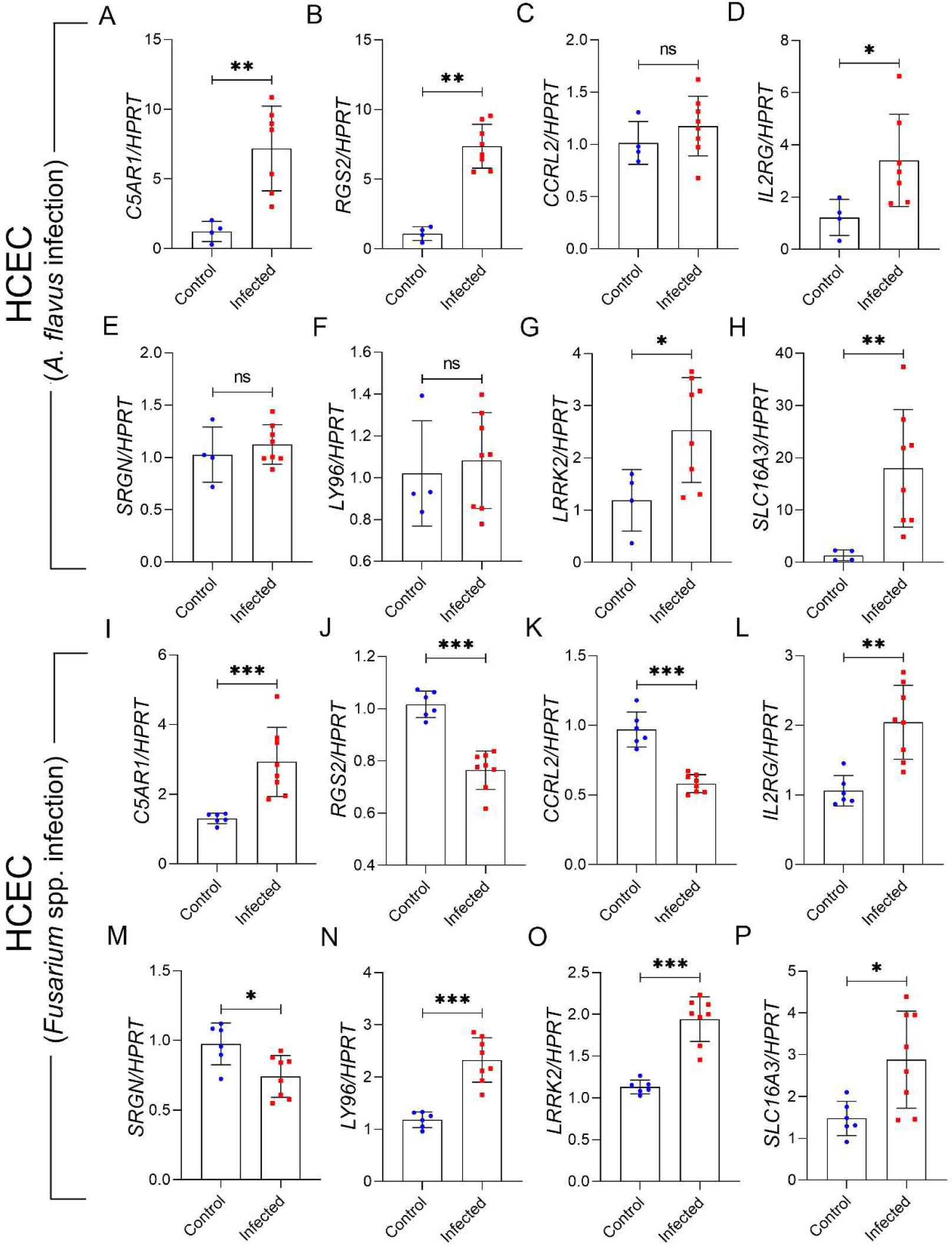
Expression of *Aspergillus flavus* keratitis-associated genes in HCECs following fungal infection. HPRT-normalized expression of candidate genes in human corneal epithelial cells (HCECs) infected with *A. flavus* spores (A-H) or *Fusarium* spores (I-P), shown as fold change relative to uninfected controls. Bars represent mean ± SD with individual data points representing biological replicates. Statistical significance was determined by Mann-Whitney U test: \**P*<.05; \*\**P*<.01; \*\*\**P*<.001; ns, nonsignificant.

Among *Fusarium*-derived candidates, LPS induced *ICOSL* and *IL17C* in both cell types, whereas robust *PDE4B* induction was restricted to differentiated THP-1 cells (Supplementary Figure 10). Fungal challenge revealed distinct pathogen- and cell-specific responses. In HCECs, *Fusarium* induced *ICOSL, IL17C,* and *CREB5,* whereas *A. flavus* induced all five genes, including *SLC26A4* and *PDE4B* (Figure 4A-4E, 4K-4O). In differentiated THP-1 cells, *Fusarium* increased *PDE4B* while reducing *SLC26A4*, whereas *A. flavus* induced *IL17C, CREB5,* and *PDE4B* (Figure 4F-4J, 4P-4T). Thus, *A. flavus* elicited a broader transcriptional response, whereas responses to *Fusarium* were more gene- and cell-context dependent.

The *A. flavus* severity/outcome-associated gene panel was next examined for transcriptional responses to inflammatory and fungal challenge. LPS elicited limited responses in HCECs and mixed, predominantly suppressive responses in differentiated THP-1 cells (Supplementary Figure 11), whereas live fungal challenge revealed marked pathogen- and cell-specific patterns. In HCECs, *A. flavus* induced *C5AR1, RGS2, IL2RG, LRRK2,* and *SLC16A3*, whereas *Fusarium* produced a mixed response, inducing *C5AR1, IL2RG, LY96, LRRK2,* and *SLC16A3* while suppressing *RGS2, CCRL2,* and *SRGN* (Figure 5). The magnitude of induction was also substantially greater following *A. flavus* infection. This pathogen-specific contrast was even more pronounced in differentiated THP-1 cells: *A. flavus* induced *RGS2, CCRL2, SRGN, MYO1F, ITGAL,* and *SLC16A3,* whereas *Fusarium* broadly suppressed the panel, with *SLC16A3* remaining unchanged (Supplementary Figure 12). Thus, *A. flavus* elicited a predominantly inductive response, whereas the *Fusarium* response was strongly cell-context dependent. This pattern was concordant with the broader expression of these genes in *A. flavus* patient corneas and their relatively limited expression in healed *Fusarium* keratitis (Supplementary Figure 9), further linking pathogen-specific experimental responses with the etiology-associated host-response patterns observed in patients.

## Discussion

In this cohort study, we identified corneal gene-expression signatures associated with disease severity and treatment response across FK etiologies. To our knowledge, this is the first transcriptomic analysis of clinically stratified corneal samples at presentation across etiologies, linking host-response signatures to patient-level disease parameters. Previous work from our group showed *Fusarium* spp. to be most common but *Aspergillus flavus* associated with greater severity ^5,14,16^. Extending these observations beyond prevalence-related disease burden, our retrospective analysis compared etiology-related disease trajectories. While the poorer clinical course of *A. flavus* remained consistent, *Fusarium* emerged with the most favourable healing with preserved vision, even compared with the more frequently healed culture-negative group. These findings informed our severity- and outcome-stratified transcriptomic analysis, addressing a critical gap in linking early host responses to subsequent disease course. Together, the clinical and molecular findings indicate that etiology-related differences in disease course are, at least in part, linked to distinct corneal host-response programs.

The corneal host-response gene expression programs identified in this study comprise *ICOSL, IL-17C, SLC26A4,* and a coordinated innate inflammatory gene set. *ICOSL,* a key regulator of T-cell responses ^17–20^, is increasingly recognized for its role at epithelial–adaptive immune interfaces, including immune regulation by regulatory T cells ^24–27^. Its expression in corneal tissue, demonstrated here, identifies the *ICOS–ICOSL* axis as a previously unrecognized component of corneal immune regulation. Association of higher corneal *ICOSL* expression with a favourable response in *Fusarium* keratitis and its strong induction in corneal epithelial cells *in vitro* indicates an epithelial contribution to this protective mechanism. *IL-17C,* an epithelial-derived cytokine known to mediate autocrine innate and proinflammatory responses, was broadly upregulated irrespective of severity, consistent with a generalized epithelial alarm response in fungal keratitis ^8,21,22^. *SLC26A4* (pendrin), an epithelial anion transporter implicated in mucosal inflammation and innate defense ^23,24^, was associated with worse outcomes in *Fusarium* keratitis, suggesting barrier dysfunction and inflammatory injury in the cornea.

The inflammatory module—including *LY96, LRRK2, SRGN, C5AR1, CCRL2, IL2RG, MYO1F, ITGAL,* and *CREB5*, along with regulatory molecules such as *PDE4B* and *RGS2*—defines a coordinated program of innate immune activation, leukocyte recruitment, and inflammatory amplification, extending previous transcriptomic and proteomic evidence of innate immune–driven inflammation in advanced FK ^8,9,11^ by resolving specific upstream and downstream components within this network. Activation of Toll-like receptor signaling, reflected by upregulation of *LY96* (MD-2) ^25,26^ and *LRRK2* ^27–29^, supports engagement of NF-κB–mediated cytokine pathways (e.g., IL-1β, TNF-α, IL-6) ^30,31^, consistent with prior reports ^10,15^. Concurrent upregulation of *CCRL2, MYO1F,* and *SRGN* indicates active neutrophil recruitment and degranulation ^32–34^, in agreement with proteomic detection of canonical neutrophil-derived proteins such as *S100A8/A9* and myeloperoxidase in tear fluid ^12,35^. Increased *C5AR1* further supports complement activation, previously demonstrated in fungal keratitis ^11,13^ indicating C5a-mediated inflammatory signalling relevant to antifungal defense ^36,37^.

Our dataset further implicates cAMP-responsive signaling through increased *PDE4B*, a cAMP-specific phosphodiesterase ^38,39^, and *CREB5*, a cAMP-responsive transcription factor ^40,41^. *PDE4B* promotes inflammatory signaling by relieving cAMP-mediated inhibition of NF-κB and *NLRP3* inflammasome pathways ^42–44^, key innate drivers of inflammation in FK. Its robust induction following *A. flavus* infection in corneal epithelial and THP-1 cells, compared with attenuated *Fusarium* responses, mirrored patient profiles and is consistent with its reported role in ocular inflammation including uveitis ^45–47^. *RGS2*, a negative regulator of GPCR signaling^48^, having implications in intraocular pressure regulation and age related corneal dystrophies ^49,50^, was preferentially upregulated in severe *A. flavus* keratitis. In vitro, *RGS2* was induced by *A. flavus* but suppressed by *Fusarium*, implying differential engagement of GPCR-mediated inflammatory pathways across etiologies.

Beyond identifying novel markers within established pathways, our findings define coordinated host-response profiles associated with fungal keratitis etiology and outcome. Elevated *ICOSL* with limited inflammatory activation was associated with favourable response to therapy, predominantly in *Fusarium* keratitis, whereas marked activation of the inflammatory module was linked to poor clinical and visual outcomes in *A. flavus*. Particularly important is culture-negative keratitis, which, despite frequent clinical healing, showed poor visual outcomes and substantial overlap with the *A. flavus* inflammatory profile. This molecular–clinical discordance provides new insight into an etiologically heterogeneous and poorly characterized group, implicating excessive host innate responses as a key driver of pathogenesis in culture-negative keratitis, highlighting the importance of host-directed therapy for better clinical outcomes, potentially in preservation of vision.

### Limitations

Our data are limited in validating healing-associated protective responses beyond *ICOSL*. While they support identification of a predictive biomarker panel for disease course, large-scale validation is required to establish their utility as biomarkers.

## Conclusion

In summary, we identified gene signatures associated with disease severity and outcomes, suggesting that corneal molecular profiling at presentation may enable early prediction and guide therapy. These insights provide a foundation for predictive biomarker panels for risk stratification and targets for host-directed immunomodulation.

## Supporting information

Supplement_BioRxive_FK_2026

## Acknowledgements

We thank Professor K. Dharmalingam, PhD (Director-Research, Aravind Medical Research Foundation), for his critical comments, discussions, and valuable suggestions, and G. Rameshkumar, MSc (Aravind Eye Hospital), for assistance with retrospective data collection and provision of clinical isolates of *Fusarium* spp. and *Aspergillus flavus*. Neither individual received compensation for their contributions. We also thank Vikram Subramanian, CEO and MD of Biokart India Pvt Ltd, for assistance with RNA-sequencing raw data processing and analysis.

## Declaration of generative AI and AI-assisted technologies in the writing process

GPT-5.5 (ChatGPT, OpenAI) was used solely for grammatical and language editing of the manuscript. The authors reviewed and approved all AI-assisted edits and take full responsibility for the manuscript content, including the text, results, data, analyses, interpretation, and conclusions.

## References

1. Khor WB, Prajna VN, Garg P, et al. The Asia Cornea Society Infectious Keratitis Study: A Prospective Multicenter Study of Infectious Keratitis in Asia. American Journal of Ophthalmology. 2018;195:161–170. doi:10.1016/j.ajo.2018.07.040

2. Ung L, Bispo PJ, Shanbhag SS, Gilmore MS, Chodosh J. The persistent dilemma of microbial keratitis: Global burden, diagnosis, and antimicrobial resistance. Survey of ophthalmology. 2019;64(3):255–271.

3. Gopinathan U, Sharma S, Garg P, Rao GN. Review of epidemiological features, microbiological diagnosis and treatment outcome of microbial keratitis: Experience of over a decade. Indian Journal of Ophthalmology. 2009;57(4):273. doi:10.4103/0301-4738.53051

4. Mills B, Radhakrishnan N, Karthikeyan Rajapandian SG, Rameshkumar G, Lalitha P, Prajna NV. The role of fungi in fungal keratitis. Experimental Eye Research. 2021;202:108372. doi:10.1016/j.exer.2020.108372

5. Prajna NV, Krishnan T, Mascarenhas J, et al. The Mycotic Ulcer Treatment Trial: A Randomized Trial Comparing Natamycin vs Voriconazole. JAMA Ophthalmol. 2013;131(4):422–429. doi:10.1001/jamaophthalmol.2013.1497

6. Sun CQ, Prajna NV, Krishnan T, et al. Effect of pretreatment with antifungal agents on clinical outcomes in fungal keratitis. Clinical Exper Ophthalmology. 2016;44(9):763–767. doi:10.1111/ceo.12794

7. Prajna NV, Lalitha P, Krishnan T, et al. Patterns of Antifungal Resistance in Adult Patients With Fungal Keratitis in South India: A Post Hoc Analysis of 3 Randomized Clinical Trials. JAMA Ophthalmol. 2022;140(2):179–184. doi:10.1001/jamaophthalmol.2021.5765

8. Lapp T, Kammrath Betancor P, Schlunck G, et al. Transcriptional profiling specifies the pathogen-specific human host response to infectious keratitis. Front Cell Infect Microbiol. 2024;13. doi:10.3389/fcimb.2023.1285676

9. Chidambaram JD, Kannambath S, Srikanthi P, et al. Persistence of Innate Immune Pathways in Late Stage Human Bacterial and Fungal Keratitis: Results from a Comparative Transcriptome Analysis. Front Cell Infect Microbiol. 2017;7. doi:10.3389/fcimb.2017.00193

10. Dinesh S, Praveenkumar R, Lalitha P, Prajna NV, Dharmalingam K, Devarajan B. Dysregulation of human corneal miRNAs and their cognate targets in *Fusarium* spp. keratitis and its implication in disease progression. The Ocular Surface. 2026;40:206–215. doi:10.1016/j.jtos.2026.02.012

11. Alenezi H, Parnell G, Schibeci S, et al. Ocular surface immune transcriptome and tear cytokines in corneal infection patients. Front Cell Infect Microbiol. 2024;14. doi:10.3389/fcimb.2024.1346821

12. Kandhavelu J, Demonte NL, Namperumalsamy VP, et al. *Aspergillus flavus* induced alterations in tear protein profile reveal pathogen-induced host response to fungal infection. Journal of Proteomics. 2017;152:13–21. doi:10.1016/j.jprot.2016.10.009

13. Shait Mohammed MR, Krishnan S, Amrathlal RS, et al. Local Activation of the Alternative Pathway of Complement System in Mycotic Keratitis Patient Tear. Front Cell Infect Microbiol. 2020;10. doi:10.3389/fcimb.2020.00205

14. Lalitha P, Prajna NV, Kabra A, Mahadevan K, Srinivasan M. Risk Factors for Treatment Outcome in Fungal Keratitis. Ophthalmology. 2006;113(4):526–530. doi:10.1016/j.ophtha.2005.10.063

15. Dinesh S, Prajna L, Venkatesh PN, Dharmalingam K, Devarajan B. Fusarium spp. and Aspergillus flavus infection induces pathogen-specific and pathogen-independent host immune response in patients with fungal keratitis. Front Cell Infect Microbiol. 2025;15. doi:10.3389/fcimb.2025.1560628

16. Lalitha P, Prajna NV, Oldenburg CE, et al. Organism, Minimum Inhibitory Concentration, and Outcome in a Fungal Corneal Ulcer Clinical Trial. Cornea. 2012;31(6):662. doi:10.1097/ICO.0b013e31823f8ae0

17. Dong C, Juedes AE, Temann UA, et al. ICOS co-stimulatory receptor is essential for T-cell activation and function. Nature. 2001;409(6816):97–101. doi:10.1038/35051100

18. Mak TW, Shahinian A, Yoshinaga SK, et al. Costimulation through the inducible costimulator ligand is essential for both T helper and B cell functions in T cell–dependent B cell responses. Nat Immunol. 2003;4(8):765–772. doi:10.1038/ni947

19. Greenwald RJ, Freeman GJ, Sharpe AH. THE B7 FAMILY REVISITED. Annual Review of Immunology. 2005;23(Volume 23, 2005):515–548. doi:10.1146/annurev.immunol.23.021704.115611

20. Akbari O, Stock P, Meyer EH, et al. ICOS/ICOSL Interaction Is Required for CD4+ Invariant NKT Cell Function and Homeostatic Survival1. J Immunol. 2008;180(8):5448–5456. doi:10.4049/jimmunol.180.8.5448

21. Ramirez-Carrozzi V, Sambandam A, Luis E, et al. IL-17C regulates the innate immune function of epithelial cells in an autocrine manner. Nat Immunol. 2011;12(12):1159–1166. doi:10.1038/ni.2156

22. Song X, Zhu S, Shi P, et al. IL-17RE is the functional receptor for IL-17C and mediates mucosal immunity to infection with intestinal pathogens. Nat Immunol. 2011;12(12):1151–1158. doi:10.1038/ni.2155

23. Jia CE, Jiang D, Dai H, Xiao F, Wang C. Pendrin, an anion exchanger on lung epithelial cells, could be a novel target for lipopolysaccharide-induced acute lung injury mice. Am J Transl Res. 2016;8(2):981–992.

24. Vanoni S, Scantamburlo G, Dossena S, Paulmichl M, Nofziger C. Interleukin-Mediated Pendrin Transcriptional Regulation in Airway and Esophageal Epithelia. International Journal of Molecular Sciences. 2019;20(3):731. doi:10.3390/ijms20030731

25. Fu Y, Kim H, Lee DS, et al. Structural insight into TLR4/MD-2 activation by synthetic LPS mimetics with distinct binding modes. Nat Commun. 2025;16(1):4164. doi:10.1038/s41467-025-59550-3

26. Bai Y, Min R, Chen P, et al. Disulfiram blocks inflammatory TLR4 signaling by targeting MD-2. Proceedings of the National Academy of Sciences. 2023;120(31):e2306399120. doi:10.1073/pnas.2306399120

27. Ho DH, Nam D, Seo M, Park SW, Seol W, Son I. LRRK2 Inhibition Mitigates the Neuroinflammation Caused by TLR2-Specific α-Synuclein and Alleviates Neuroinflammation-Derived Dopaminergic Neuronal Loss. Cells. 2022;11(5):861. doi:10.3390/cells11050861

28. Oun A, Hoeksema E, Soliman A, et al. Characterization of Lipopolysaccharide Effects on LRRK2 Signaling in RAW Macrophages. International Journal of Molecular Sciences. 2023;24(2):1644. doi:10.3390/ijms24021644

29. Wallings RL, Herrick MK, Tansey MG. LRRK2 at the Interface Between Peripheral and Central Immune Function in Parkinson’s. Front Neurosci. 2020;14. doi:10.3389/fnins.2020.00443

30. Kubo M, Nagashima R, Kurihara M, et al. Leucine-Rich Repeat Kinase 2 Controls Inflammatory Cytokines Production through NF-κB Phosphorylation and Antigen Presentation in Bone Marrow-Derived Dendritic Cells. International Journal of Molecular Sciences. 2020;21(5):1890. doi:10.3390/ijms21051890

31. Cao J, Yang Z. FOXA2 inhibits the TLR4/NF-κB signaling pathway and alleviates inflammatory activation of macrophages in rheumatoid arthritis by repressing LY96 transcription. Cytokine. 2024;184:156796. doi:10.1016/j.cyto.2024.156796

32. Del Prete A, Martínez-Muñoz L, Mazzon C, et al. The atypical receptor CCRL2 is required for CXCR2-dependent neutrophil recruitment and tissue damage. Blood. 2017;130(10):1223–1234. doi:10.1182/blood-2017-04-777680

33. Salvermoser M, Pick R, Weckbach LT, et al. Myosin 1f is specifically required for neutrophil migration in 3D environments during acute inflammation. Blood. 2018;131(17):1887–1898. doi:10.1182/blood-2017-10-811851

34. Kolset SO, Pejler G. Serglycin: A Structural and Functional Chameleon with Wide Impact on Immune Cells. J Immunol. 2011;187(10):4927–4933. doi:10.4049/jimmunol.1100806

35. Ananthi S, Venkatesh Prajna N, Lalitha P, Valarnila M, Dharmalingam K. Pathogen induced changes in the protein profile of human tears from Fusarium keratitis patients. PLoS One. 2013;8(1):e53018.

36. Desai JV, Kumar D, Freiwald T, et al. C5a-licensed phagocytes drive sterilizing immunity during systemic fungal infection. Cell. 2023;186(13):2802–2822.e22. doi:10.1016/j.cell.2023.04.031

37. Desai JV, Lionakis MS. C5-C5aR1-mediated immune responses during fungal infection: Clinical and translational implications. Clin Transl Med. 2023;13(9):e1424. doi:10.1002/ctm2.1424

38. Zhang Y, Yang Y, Liang H, et al. Nobiletin, as a Novel PDE4B Inhibitor, Alleviates Asthma Symptoms by Activating the cAMP-PKA-CREB Signaling Pathway. International Journal of Molecular Sciences. 2024;25(19):10406. doi:10.3390/ijms251910406

39. Jin SLC, Conti M. Induction of the cyclic nucleotide phosphodiesterase PDE4B is essential for LPS-activated TNF-α responses. Proceedings of the National Academy of Sciences. 2002;99(11):7628–7633. doi:10.1073/pnas.122041599

40. Wang S, Qiu J, Liu L, et al. CREB5 promotes invasiveness and metastasis in colorectal cancer by directly activating MET. J Exp Clin Cancer Res. 2020;39(1):168. doi:10.1186/s13046-020-01673-0

41. Long X, Li Y, Qiu S, Liu J, He L, Peng Y. MiR-582-5p/miR-590-5p targeted CREB1/CREB5–NF-κB signaling and caused opioid-induced immunosuppression in human monocytes. Transl Psychiatry. 2016;6(3):e757–e757. doi:10.1038/tp.2016.4

42. Wang Y, Sun D, Dilixiati N, Wang D, Song Y, Ye Q. Nerandomilast, a PDE4B inhibitor, alleviates silica-induced lung inflammation and fibrosis by inhibition of NLRP3 inflammasome and TGF-β/Smad signalling. British Journal of Pharmacology. 2026;183(7):1445–1464. doi:10.1111/bph.70240

43. Dhar R, Rana MN, Zhang L, et al. Phosphodiesterase 4B is required for NLRP3 inflammasome activation by positive feedback with Nrf2 in the early phase of LPS-induced acute lung injury. Free Radical Biology and Medicine. 2021;176:378–391. doi:10.1016/j.freeradbiomed.2021.10.007

44. Dhar R, Li Y, Hu Z, et al. PDE4B Modulates Phosphorylation of p65 (Ser468) via cAMP/PKA in Acute Lung Injury. Lung. 2025;203(1):33. doi:10.1007/s00408-025-00787-6

45. Kondapaneni LP, Arora M, Scott EM, Kumar MNVR, Ganugula R. Establishing PDE4 as a Novel Target of Urolithin-A in Mitigating LPS-induced Inflammation in Retinal Pigmented Epithelium Cells. Pharm Res. 2025;42(12):2173–2184. doi:10.1007/s11095-025-03933-0

46. Takeno M. Positioning of apremilast in treatment of Behçet’s disease. Mod Rheumatol. 2020;30(2):219–224. doi:10.1080/14397595.2019.1696504

47. Chen B, Huang Z, Chen J, et al. Effect of Dipyridamole on Experimental Autoimmune Uveitis: Reprogrammed Immune Cell Landscape and Reduced Th17 Pathogenicity. Invest Ophthalmol Vis Sci. 2025;66(15):16. doi:10.1167/iovs.66.15.16

48. Joshi JC, Joshi B, Zhang C, et al. RGS2 is an innate immune checkpoint for suppressing Gαq-mediated IFNγ generation and lung injury. iScience. 2025;28(2). doi:10.1016/j.isci.2025.111878

49. Inoue-Mochita M, Inoue T, Epstein DL, Blumer KJ, Rao PV. RGS2-deficient mice exhibit decreased intraocular pressure and increased retinal ganglion cell survival. Mol Vis. 2009;15:495–504.

50. Briere A, Vo P, Yang B, et al. Systematic Ocular Phenotyping of Knockout Mouse Lines Identifies Genes Associated With Age-Related Corneal Dystrophies. Invest Ophthalmol Vis Sci. 2025;66(5):7. doi:10.1167/iovs.66.5.7

