## Supplement_BioRxive_FK_2026 for "Early Corneal Host-Response Signatures Associate with Disease Severity and Etiology-Related Clinical Outcomes in Fungal Keratitis"

**Content:**

**Supplementary Methods.**

**Supplementary Table 1.** Oligonucleotide sequences used for RT-qPCR.

**Supplementary Table 2.** Demographic and clinical characteristics of the patient cohort in retrospective study.

**Supplementary Table 3.** Comparison of clinical presentation and treatment outcomes across etiologies of keratitis.

**Supplementary Table 4.** Comparison of clinical presentation and treatment outcomes across etiologies in a 3-month consecutive keratitis cohort.

**Supplementary Table 5.** Comparison of treatment and visual outcomes across etiologies of fungal keratitis.

**Supplementary Table 6.** Comparison of ulcer location across etiologies among healed patients.

**Supplementary Table 7.** Candidate healing-associated signatures in *Fusarium* keratitis.

**Supplementary Table 8.** Candidate unhealing-associated signatures in *Fusarium* keratitis.

**Supplementary Table 9.** Candidate severity/outcome-associated signatures in *A. flavus* keratitis

**Supplementary Table 10.** Demographic and clinical characteristics of the validation study cohort

**Supplementary Table 11.** Area under receiver operating characteristic analysis of severity-associated signatures.

**Supplementary Table 12. Gene expression profiles demonstrate outcome-associated clustering patterns in *A. flavus* keratitis.**

**Supplementary Table 13**. Gene expression profiles demonstrate outcome-associated clustering patterns in *Fusarium* and *A. flavus* keratitis.

**Supplementary Table 14.** Gene expression profiles demonstrate outcome-associated clustering patterns in microbial keratitis.

**Supplementary Table 15.** Gene expression–based prediction of clinical severity and outcomes in a masked keratitis cohort.

**Supplementary Table 16.** Candidate framework for interpretation of presentation-stage host-response profiles

**Supplementary Figure 1.** Comparison of clinical presentation and outcomes across etiologies of keratitis.

**Supplementary Figure 2.** Transcriptomic profiling and identification of host-response gene signatures associated with clinical trajectory in *Fusarium* Keratitis.

**Supplementary Figure 3.** Unsupervised hierarchical clustering of the Fu_151 gene subset in *Fusarium* keratitis showing potential healing-associated signature.

**Supplementary Figure 4.** Unsupervised hierarchical clustering of the Fu_195 gene subset in *Fusarium* keratitis showing potential unhealing-associated signature.

**Supplementary Figure 5.** Unsupervised hierarchical clustering of the Afl_287 gene subset in Aspergillus flavus showing potential severity/outcome-associated signature.

**Supplementary Figure 6.** Comparative transcriptomic analysis and identification of shared host-response gene signatures across keratitis etiologies.

**Supplementary Figure 7.** Validation of gene expression in *Fusarium* and *A. flavus* keratitis patient samples.

**Supplementary Figure 8.** Comparison of outcome-associated gene expression signatures of *Fusarium* keratitis with healed groups of other etiologies.

**Supplementary Figure 9.** Comparison of severity/outcome-associated gene expression signatures of *A.flavus* keratitis with healed groups of other etiologies.

**Supplementary Figure 10.** Effect of LPS stimulation on the outcome-associated candidate genes of *Fusarium* keratitis in cell models.

**Supplementary Figure 11.** Effect of LPS stimulation on the severity/outcome-associated candidate genes of *A.flavus* keratitis in cell models.

**Supplementary Figure 12.** Effect of fungal spore co-culture on the severity/outcome-associated candidate genes of *A.flavus* keratitis in cell models.

**Supplementary Figure 13.** Gene Ontology (GO) enrichment analysis of DEGs from *Fusarium* and *A. flavus keratitis* RNA-seq datasets.

**Supplementary Methods.**

**Retrospective study participants and design:**

Between January 2021 and December 2024, a total of 483 and 1255 patients were treated for *Aspergillus flavus* and *Fusarium* *spp.* respectively, at the Cornea Clinic, Aravind Eye Hospital, Madurai. During the same period 4189 patients were reported culture negative by standard microbiological analysis. Among these, patients who completed treatment were included in the retrospective analysis, comprising 86.9% of *A. flavus*, 38.3% of *Fusarium* *spp.*, and 9.6% of culture-negative keratitis cases. A summary of the cohort demographics is provided in **Supplementary Table 2.**

**Prospective study participants, design and Sample Collection:**

The study adhered to the tenets of the Declaration of Helsinki and was approved by the Institutional Ethics Committee. Written informed consent was obtained from all participants. Patients aged ≥13 years presenting at their first visit with clinically presumed fungal keratitis and undergoing corneal ulcer scraping for microbiological analysis were eligible. All samples were collected under topical anesthesia with 0.5% proparacaine. Corneal scrapings were obtained from the epithelium at the ulcer margin using a sterile blade or spatula. The scraped area was then gently mopped with a sterile polyester-tipped swab (HiMedia Laboratories) to collect residual material. Both the scraping instrument and the swab tip were immediately placed into cryovials containing 300 µL of DNA/RNA Shield (Zymo Research, Irvine, CA), maintained on ice, and stored at −80 °C until RNA extraction.

All participants were followed throughout treatment until the final visit to determine clinical outcomes. For RNA sequencing, samples were grouped based on the infecting pathogen (e.g., *Fusarium spp*, *Aspergillus flavus*, and culture-negative cases). Samples were further categorized by baseline clinical severity (ulcer size and depth) as non-severe or severe, and subsequently stratified by treatment outcome as healed (resolution of infection with corneal healing) or unhealed (failure of medical therapy requiring therapeutic penetrating keratoplasty/TPK). For RT-qPCR validation, samples were initially classified only by pathogen (*Fusarium spp*, *Aspergillus flavus*, *Curvularia* *spp., Pseudomonas aeruginosa*, culture negative) and molecular analyses were performed prior to the knowledge of clinical outcomes or any clinical parameters. Following analysis, results were stratified according to final treatment outcomes. In a subset of six patients, RT-qPCR analysis was performed prior to pathogen identification, enabling prediction of disease severity and clinical outcome in a fully masked setting.

For the validation cohort, clinical data, including baseline severity (ulcer size and depth), uncorrected visual acuity, post-keratoplasty complications, and final visual acuity- were collected after completion of molecular analyses and used to assess associations with gene expression data. Patient demographics for the validation cohort are provided in **Supplementary Table 10**. Details of participants contributing RNA sequencing data are available in the NCBI Sequence Read Archive (PRJNA1406027).

**Cell culture, LPS stimulation and infection with *Fusarium spp.* or *Aspergillus flavus* spores**

HCECs (RCB2280) were cultured in DMEM/F12 (Gibco, Thermo Fisher Scientific) supplemented with 5% fetal bovine serum (FBS; Gibco), 10 ng/mL recombinant human epidermal growth factor (Gibco), 5 µg/mL human recombinant insulin (Gibco), 0.5% dimethyl sulfoxide (Thermo Fisher Scientific), and 1% penicillin/streptomycin at 37 °C in 5% CO₂. THP-1 cells were maintained in RPMI-1640 (Gibco, Thermo Fisher Scientific) supplemented with 10% FBS and 1% penicillin/streptomycin under the same conditions.For macrophage differentiation, THP-1 cells were seeded at 0.5 × 10⁶ cells/mL and treated with phorbol 12-myristate 13-acetate (PMA; 10 ng/mL; Sigma-Aldrich) for 24 hours, followed by incubation in fresh serum-containing medium for an additional 24 hours.

For stimulation experiments, HCECs and differentiated THP-1 cells (approximately 50% confluence) were serum-starved for 3 hours and treated with lipopolysaccharide (LPS; 100 ng/mL; Sigma-Aldrich) for 5, 16, or 24 hours. For in vitro infection assays, clinical isolates of Aspergillus flavus and Fusarium species were obtained from the Microbiology Laboratory, Aravind Eye Hospital (Madurai, India). Fungi were cultured on potato dextrose agar at 30 °C for 5–7 days. Spores were harvested using 0.05% Tween 20, filtered, and enumerated using a Neubauer chamber. Spores were resuspended in serum-free medium, allowed to swell for 1 hour, and added to cells at a multiplicity of infection of 30:1 (spores to host cells) for 8 hours at 37 °C in 5% CO₂.

**RNA extraction:**

Stored corneal swabs in cryovials and cell lysates from in vitro LPS stimulation and infection experiments were thawed on ice and processed in batches of no more than six to preserve nucleic acid integrity. Swabs were transferred to 2-mL tubes containing 800 µL of TRIzol reagent (Invitrogen, Thermo Fisher Scientific, Waltham, MA) and thoroughly mixed by agitation and pipetting. Residual Zymo buffer was collected, transferred to 1.5-mL tubes, and centrifuged at 3000 rpm for 10 minutes at 4 °C to pellet remaining cells. After removal of the supernatant, leaving approximately 200 µL, an equal volume of TRIzol was added, and the mixture was combined with the corresponding swab-derived samples.

For both patient-derived and cell-derived samples, chloroform (200 µL) was added to initiate phase separation, followed by vortexing in three 10-second pulses, incubation on ice for 5 minutes, and centrifugation at 12,000 rpm for 15 minutes at 4 °C. The aqueous phase was transferred to fresh 1.5-mL tubes, and RNA was precipitated with an equal volume of isopropanol. For patient-derived samples, 10 µg glycogen (Thermo Fisher Scientific) was added as a carrier. After incubation at 4 °C for 1 hour, samples were centrifuged at 12,000 rpm for 10 minutes. The RNA pellet was washed with 1 mL of 75% ethanol, centrifuged at 7,500 rpm for 5 minutes, air-dried, and resuspended in RNase-free water.

**RNA sequencing, data processing and analysis:**

RNA sequencing was conducted using pooled RNA from multiple patient samples (n = 2–4) within each defined group, yielding 1.5–2 µg total RNA per pool, with at least three pooled biological replicates analyzed per group. RNA concentration was measured using a NanoDrop ND-LITE, and integrity was assessed with an Agilent Bioanalyzer 2100; only samples with 260/280 ratios between 1.8 and 2.2 and RNA integrity numbers (RIN) >7 were advanced for library preparation using the NEBNext® Ultra™ Directional RNA Library Prep Kit (Illumina®). Sequencing was carried out on an Illumina HiSeq 4000 platform. Raw sequencing reads were processed by quality trimming with the FASTX toolkit and aligned to the GRCh37 human reference genome using STAR. Gene-level read counts were obtained with FeatureCounts and subsequently normalized using the Limma-Voom method. Differential gene expression analysis was performed using iGEAK, with significance defined as an adjusted p-value <0.05. Principal component analysis (PCA) was performed in R using normalized expression data (Limma-Voom), and visualized using custom R scripts to assess sample clustering and variability across groups.

Differentially expressed gene (DEG) lists across comparisons were analyzed to identify shared and unique patterns. Overlaps were determined using Microsoft Access and visualized as Venn diagrams with Molbiotools, enabling identification of gene subsets with potential transcriptomic signatures. These subsets were further analyzed for Gene Ontology enrichment using g: Profiler to define associated biological processes. Signature robustness was assessed by unsupervised hierarchical clustering of normalized expression data (Limma-Voom), visualized as heatmaps using custom R scripts, demonstrating clear group-wise segregation of samples. Volcano plots were generated using VolcaNoseR, where top genes were prioritized based on a composite ranking integrating log2 fold change and –log10 adjusted p-value.

**Identification of transcriptomic signatures of interest**

***Fusarium* keratitis**

Transcriptomic signatures associated with disease outcome were systematically derived using subgroup-specific differential expression analyses. All four clinical subgroups (non-severe healed, non-severe unhealed, severe healed, and severe unhealed) were included. Initial comparisons identified 1,379 DEGs (Fu_1379) shared across all disease subgroups relative to controls (**Supplementary Figure 2B**) and 983 DEGs (Fu_983) common to the severe-healed and unhealed groups relative to the non-severe healed group (**Supplementary Figure 2C**). Minimal overlap (63 genes, Fu_63) between these sets suggested the absence of a major common subset of genes with healed- and unhealed-associated expression profiles that are distinguishable from each other and from controls (**Supplementary Figure 2D**). However, volcano plot–based prioritization of the 983- and 63-gene subsets identified candidate severity- and outcome-associated genes (**Supplementary Figure 2G-2J**), which were not validated in this study, with the exception of PDE4B and CREB5.

To identify distinct subsets of genes associated with healed and unhealed outcomes, refined overlap analyses were performed. A healed-specific signature was defined by genes differentially expressed in the non-severe healed group relative to both the controls and the unhealed groups, excluding DEGs identified in the non-severe healed versus severe-healed comparison, yielding a 151-gene subset (Fu_151) (**Figure 1C and 1D**). In parallel, an unhealed-specific signature (195 genes, Fu_195) was derived from DEGs shared between the non-severe and severe unhealed groups relative to the non-severe healed group, excluding DEGs identified in the control versus non-severe healed and non-severe healed versus severe-healed comparisons (**Figure 1C and 1D**). Both signatures were validated by unsupervised hierarchical clustering, demonstrating distinct segregation of sample groups and outcome-associated expression profiles (**Supplementary Figure 3 and 4**).

Functional enrichment analysis of the combined 346 genes revealed pathways consistent with infection- and inflammation-associated responses (**Supplementary Figure 13**). Candidate genes from the 151- and 195-gene subsets were selected based on fold change, statistical significance, and representation within outcome-specific subsets (**Figure 1G and 1H and Supplementary Figure 2E and 2F**). Potential healing-associated candidates were included among the 102 of the 151 DEGs that were uniquely altered in the non-severe healed group (not significantly different in the severe-healed group relative to the non-severe healed) (**Figure 1E**) and not shared with DEGs in the unhealed groups relative to the control, with the exception of LILRB4. LILRB4 was the only candidate showing downregulation in healed samples and upregulation in unhealed samples relative to the control. Conversely, potential unhealed-associated candidates were included among the 71 DEGs that were shared between the non-severe healed versus unhealed comparisons and the control versus unhealed comparisons (**Figure 1F**).

***Aspergillus flavus* keratitis**

Transcriptomic analysis was performed across two clinical subgroups (non-severe healed and severe unhealed), where severity- and outcome-associated signatures were expected to overlap. DEG comparisons identified 287 genes shared between both disease groups relative to controls and in direct subgroup comparisons (**Figure 1I and 1J**). Functional enrichment of these genes highlighted biological processes associated with heightened immune signaling and inflammatory responses (**Supplementary Figure 13**).

Clustering analysis of the 287-gene subset demonstrated differential expression between healed and unhealed groups relative to controls as well as similar expression profiles between *A. flavus* keratitis and culture-negative non-severe healed samples (**Supplementary Figure 5**). Volcano plot–based prioritization of the 287-gene subsets identified candidate severity- and outcome-associated genes (**Figure 1K**). Thus, *A. flavus* keratitis dataset revealed a shared gene subset with differential expression pattern associated with severity and/or clinical outcome.

**Culture-negative keratitis and cross-disease comparisons**

In culture-negative keratitis, three subgroups (non-severe healed, severe healed, and severe unhealed) were analyzed. Initial DEG comparisons identified 1057 genes (C-ve_1057) shared across all subgroups relative to controls (**Supplementary Figure 6B**) and 1081 genes (C-ve_1081) distinguishing severe (healed and unhealed) from non-severe healed samples (**Supplementary Figure 6A**), with limited overlap (109 genes, C-ve_109) (**Supplementary Figure 6C**). These overlapping genes included several candidates identified from the *A. flavus* dataset, with volcano plot analysis confirming their prominence among the top-ranked DEGs (**Supplementary Figure 6D**).

Comparative analyses between culture-negative and *A. flavus* keratitis revealed substantial overlap, including 752 shared DEGs (C-ve_Afl_752) in non-severe healed comparisons, encompassing most *A. flavus*-derived candidates (**Supplementary Figure 6E**). Further prioritization showed strong concordance in top-ranked genes across both datasets, indicating similar underlying transcriptomic profiles (**Supplementary Figure 6F and 6G**).

Next, we compared the C-ve_Afl_752 DEGs with DEGs from *Fusarium* keratitis unhealed groups (non-severe and severe) relative to the non-severe healed group, identifying 152 DEGs common to all three keratitis types, including PDE4B, CREB5, and SRGN (**Supplementary Figure 6H**). Among A. flavus–derived candidates, IFITM2, IL2RG, and LRRK2 were also differentially expressed in severe unhealed *Fusarium* keratitis, whereas the remaining candidates shared with culture-negative keratitis were not differentially expressed in *Fusarium* unhealed samples (**Supplementary Figure 6H**). In contrast, Fusarium-derived candidates (ICOSL, SLC26A4, and IL-17C) did not differentiate healed and unhealed samples in either A. flavus or culture-negative keratitis (**Supplementary Figure 6H**). Finally, comparison of non-severe healed groups across datasets identified 1,714 overlapping DEGs, including six candidates from the A. flavus analysis (**Supplementary Figure 6I**). Together, these findings indicate that selected candidates show consistent differential expression in *Fusarium* keratitis, with partial conservation across A. flavus and culture-negative keratitis.

**qPCR Analysis**

Transcript expression of individual genes was quantified using RT-qPCR. Reactions were performed on a QuantStudio Real-Time PCR system (Applied Biosystems, Waltham, MA, USA). cDNA was synthesized from 500 ng of total RNA using the QuantiTect Reverse Transcription Kit (Qiagen, Hilden, Germany) in a 10 µL reaction, according to the manufacturer’s instructions, and subsequently diluted 1:1 with nuclease-free water to a final volume of 20 µL. For qPCR, 1 µL of cDNA was added to a 9 µL reaction mixture containing QuantiNova SYBR Green Master Mix and 0.25 µM each of forward and reverse primers. The thermal cycling conditions were as follows: initial polymerase activation and denaturation at 95 °C for 10 minutes, followed by 40 cycles of 95 °C for 15 seconds and 60 °C for 1 minute. All samples were run in duplicate. Data were acquired using QuantStudio™ Design and Analysis Software (v1.4.1). Gene expression levels were normalized to the housekeeping gene *HPRT*. Relative fold changes in gene expression were calculated using the 2^−ΔΔCt method, with healthy cadaveric controls used as the reference for patient samples and uninfected or unstimulated controls used for infected or LPS-stimulated cells. Statistical significance was defined as *P* < 0.05. The primer sequences used for qPCR are listed in **Supplementary Table 9.**

**Statistical Analysis**

Baseline demographic characteristics and systemic comorbidities were compared across keratitis types using the χ² test for the retrospective cohort and Fisher’s exact test for the prospective cohort. Continuous demographic variables in both cohorts were analyzed using one-way ANOVA. In the retrospective cohort, overall comparisons of disease severity at presentation and clinical outcomes between keratitis types were performed using the χ² test, whereas Fisher’s exact test was used for the smaller 3-month consecutive cohort. Pairwise comparisons, including severity at presentation, clinical outcomes, visual outcomes, and ulcer location, were assessed using odds ratios (ORs) with 95% confidence intervals (CIs) derived from univariate, unadjusted logistic regression models. ­­For patient samples, relative mRNA expression levels among healthy cadaveric controls, healed, and unhealed groups were compared using the Kruskal–Wallis test followed by Dunn’s multiple comparison test. In vitro two-group comparisons (stimulated or infected cells vs. controls) were analyzed using the Mann–Whitney U test, while multi-time-point analyses for lipopolysaccharide (LPS) stimulation were conducted using two-way ANOVA with Sidak’s multiple comparisons test. The prognostic performance of individual gene expression levels (fold expression) was evaluated using area under the receiver operating characteristic (AUROC) curve analysis. Associations among normalized gene expression levels, as well as their relationships with ulcer size, were evaluated using Spearman rank correlation and correlation matrices. All tests were two-sided, with statistical significance set at P < 0.05. Analyses were performed using Stata version 17 (StataCorp, Texas, USA) and GraphPad Prism version 8.0.2 (2019).

| **Gene** |  | **Primer Sequence** | **Product size** | **Citations** |
| --- | --- | --- | --- | --- |
| *LY96* | Forward | 5'-GAAGCAGTATTGGGTCTGCAA-3' | 209 bps | Wu *et al.,* 2024 |
|  | Reverse | 5'-TTGGAAGATTCATGGTGTTGACA-3' |  |  |
| *IFITM2* | Forward | 5'-CATCCCGGTAACCCGATCAC-3' | 274 bps | Xu *et al.,* 2021 |
|  | Reverse | 5'-CACGGAGTACGCGAATGCTA-3' |  |  |
| *SLC16A3* | Forward | 5'-AGTGTGCGTGGTGTGGTCAG-3' | 110 bps | Cai *et al.,* 2025 |
|  | Reverse | 5'-CGAGGGCAGGTGGGTGTTAG-3' |  |  |
| *CREB5* | Forward | 5'-GCTTTGGTGCTTTTCTCCAG-3' | 245bps | Huttunen *et al.,* 2007 |
|  | Reverse | 5'-GGTGACACCACAGCACAAAC-3' |  |  |
| *LRRK2* | Forward | 5'-TGAATGCCACCGAGGAATCTG-3' | 260 bps | Miklossy *et al.,* 2006 |
|  | Reverse | 5'-GTGAACTGCGTGAGGAAGCT-3' |  |  |
| *ITGAL* | Forward | 5'-TGCTTATCATCATCACGGATGG-3' | 112 bps | Wang *et al.,* 2023 |
|  | Reverse | 5'-CTCTCCTTGGTCTGAAAATGCT-3' |  |  |
| *IL2RG* | Forward | 5'-GTGCAGCCACTATCTATTCTCTG-3' | 187 bps | Hu *et al.,* 2024 |
|  | Reverse | 5'-GTGAAGTGTTAGGTTCTCTGGAG |  |  |
| *MYO1F* | Forward | 5'-CCCCCAAGATCCCCCTGAACC-3' | 136bps | Chi *et al.,* 2024 |
|  | Reverse | 5'-ACGCCGCTCTGCTTCACGTT-3' |  |  |
| *C5AR1* | Forward | 5' -TATGGGCACTATGATGACAAGGATACC-3' | 145 bps | Xu *et al.,* 2023 |
|  | Reverse | 5'-AAGACGACTGCAAAGATGACCAAGG-3' |  |  |
| *PDE4B* | Forward | 5'-AACGCTGGAGGAATTAGACTGG-3' | 110 bps | Raza *et al.,* 2022 |
|  | Reverse | 5'-GCTCCCGGTTCAGCATTCT-3' |  |  |
| *SRGN* | Forward | 5'-CTCAGTTCAAGGTTATCCTACGC-3' | 208 bps | Villena Toro *et al.,* 2024 |
|  | Reverse | 5'-GTCCTCAGAAAGTGGGAAGATAC-3' |  |  |
| *RGS2* | Forward | 5'-CGAGGAGAAGCGAGAAAAGA-3' | 151 bps | Smalley *et al.,* 2007 |
|  | Reverse | 5'-TTCCTCAGGAGAAGGCTTGA-3' |  |  |
| *CCRL2* | Forward | 5’TGCCGCTGTTTCCATCTGCGTA-3’ | 100 bps | Reyes *et al.,* 2019 |
|  | Reverse | 5’-ACACTTCGGTGGAATGGTCAGG-3' |  |  |
| *LILRB4* | Forward | 5'-CTGTGTCAGTCACGGAGCC-3' | 190bps | Brown *et al.,* 2009 |
|  | Reverse | 5'-GCAGGTAGTGGGAGAAGCC-3' |  |  |
| *ICOSL* | Forward | 5′-TGACATCGGAGAGAGAGACAAG-3′ | 402 bps | K *et al.,* 2022 |
|  | Reverse | 5′-ACGCTTTTGAAAGGGCCTCA-3′ |  |  |
| *IL-17C* | Forward | 5′- CCCTCAGCTACGACCCAGT - 3′ | 126 bps | Lee *et al.,* 2020 |
|  | Reverse | 5′- CTTCTGTGGATAGCGGTCCT - 3′ |  |  |
| *SLC26A4* | Forward | 5′-TGGTGGCTTGCAGATTGGAT-3′ | 99 bps | Xu *et al.,* 2017 |
|  | Reverse | 5′-AGCTGTGAGACCAGCACTTG-3′ |  |  |
| *FAM53A* | Forward | 5'-GACACGCCCATCCTTGGACTTC-3' | 124 bps | *OriGene Technologies, n.d.* |
|  | Reverse | 5'-GGAGGACAGAACCGTCTTCACT-3' |  |  |
| *PNPLA3* | Forward | 5'-AATGTCCACCAGCTCATCTC-3' | 200 bps | Yuan *et al.,* 2020 |
|  | Reverse | 5'-TCACTCACTCCTCCATCCAC-3' |  |  |

**Supplementary Table 1**

**Supplementary Table 1.** **Oligonucleotide sequences used for RT-qPCR.**

**Supplementary Table 2.**

| **Characteristic** | **Total** | ***A. flavus* keratitis**  (n=420) | ***Fusarium* keratitis**  (n=481) | **Culture -ve keratitis**  (n=403) | ***P* value** |
| --- | --- | --- | --- | --- | --- |
| Male participants, No. (%) | 813 (62.4) | 258 (61.4) | 290 (60.3) | 265 (65.8) | 0.222* |
| Female participants, No. (%) | 491 (37.7) | 162 (38.6) | 191 (39.7) | 138 (34.2) |  |
| Age, mean (SD), y | 50.07 (16.2) | 52.77 (15.2) | 46.97 (16.4) | 50.95 (16.2) | <0.001^#^ |
| Age range, y | 0 – 90 | 3 - 83 | 1 - 86 | 0 - 90 | - |
| Trauma cases, No. (%) | 731 (56.1) | 250 (59.5) | 261 (54.3) | 220 (54.6) | 0.220* |
| Diabetes, No. (%) | 249 (19.1) | 121 (28.8) | 66 (13.7) | 62 (15.4) | <0.001* |
| Hypertension, No. (%) | 171 (13.1) | 63 (15.0) | 44 (9.2) | 64 (15.9) | 0.005* |
| Cardiac, No. (%) | 23 (1.8) | 7 (1.7) | 3 (0.6) | 13 (3.2) | 0.014* |

**Supplementary Table 2. Demographic and clinical characteristics of the patient cohort in retrospective study**.

Baseline demographics and systemic comorbidities of 1304 patients with *Aspergillus flavus*, *Fusarium* spp., or culture-negative keratitis. Categorical data are presented as No. (%) and continuous variables as mean (SD). *P* values indicate differences across the 3 groups, determined using the *χ*2 test (indicated by *) or 1-way ANOVA (indicated by #). Significance defined as *P* < .05.

**Supplementary Table 3.**

| **Keratitis** | **Presentation^a^**  **(*P*<0.001)** | | **OR (95% CI)^d^** | ***P* value** | **OR (95% CI)^d^** | ***P* value** | **Outcome^b^**  **(*P*<0.001)** | | **OR (95% CI)^e^** | ***P* value** | **OR (95% CI)^e^** | ***P* value** |
| --- | --- | --- | --- | --- | --- | --- | --- | --- | --- | --- | --- | --- |
| *A. flavus* (n=420) | Non-severe (50.2%) | Severe (49.8%) | 2.63 (2.01-3.43) | <0.001 | 3.64 (2.73-4.85) | <0.001 | Healed (59.52%) | Unhealed (40.48%) | 2.41 (1.80 -3.22) | <0.001 | 6.53 (4.44-9.62) | <0.001 |
| *Fusarium* spp*.* (n=481) | Non-severe (73.6%) | Severe (26.4%) | Ref. ^c^ | - | 1.39 (1.03-1.86) | 0.03 | Healed (77.96%) | Unhealed (22.04%) | Ref. ^c^ | - | 2.72 (1.82-4.04) | <0.001 |
| Culture -ve (n=403) | Non-severe (76.7%) | Severe (23.3%) | 0.72 (0.54-0.97) | 0.03 | Ref. ^c^ | - | Healed (90.57%) | Unhealed (9.43%) | 0.37 (0.25- 0.55) | <0.001 | Ref. ^c^ | - |

**Supplementary Table 3. Comparison of clinical presentation and treatment outcomes across etiologies of keratitis.**

**Abbreviations:** CI, confidence interval; OR, odds ratio.

**Footnotes:** *a* **Presentation** indicates disease severity at the time of initial clinical evaluation based on ulcer size and depth, classified into ‘non-severe’ or ‘severe’ and compared between keratitis types by *χ*2 test, *P* value is shown. *b* **Clinical outcome** indicates treatment response as ‘healed’ (infection resolution with corneal healing) or ‘unhealed’ (non-response to medical therapy requiring TPK) and compared between keratitis types by *χ*2 test, *P* value is shown. *c* **Ref.** indicates the reference category used for odds ratio calculations for presentation and clinical outcome among keratitis of different etiologies. *d* **Odds ratios for Presentation** represent the likelihood of ‘severe’ presentation, calculated using a univariate unadjusted model. *e* **Odds ratios for clinical outcome** represent the likelihood of ‘unhealed’ outcome, calculated using a univariate unadjusted model. Significance defined as *P* < .05.

**Supplementary Table 4.**

| **Keratitis** | **Presentation^a^**  **(*P*=0.077)** | | **OR (95% CI)^d^** | ***P* Value** | **OR (95% CI)^d^** | ***P* Value** | **Outcome^b^**  **(*P*<0.001)** | | **OR (95% CI)^e^** | ***P* Value** | **OR (95% CI)^e^** | ***P* Value** |
| --- | --- | --- | --- | --- | --- | --- | --- | --- | --- | --- | --- | --- |
| *A. flavus* (n=14) | Non-severe (50.0%) | Severe (50.0%) | 3.43 (1.09-10.80) | 0.035 | 3.49 (1.17-10.43) | 0.025 | Healed (35.7%) | Unhealed (64.3%) | 7.86 (2.36-26.15) | 0.001 | 17.06 (5.23-55.63) | <0.001 |
| *Fusarium spp.* (n=102) | Non-severe (77.4%) | Severe (22.6%) | Ref. ^c^ | - | 1.02 (0.58-1.78) | 0.956 | Healed (81.4%) | Unhealed (18.6%) | Ref. ^c^ | - | 2.17 (1.11-4.24) | 0.024 |
| Culture -ve (n=220) | Non-severe (77.7%) | Severe (22.3%) | 0.98 (0.56-1.73) | 0.956 | Ref. ^c^ | - | Healed (90.5%) | Unhealed (9.55%) | 0.46 (0.24-0.90) | 0.024 | Ref. ^c^ | - |

**Supplementary Table 4. Comparison of clinical presentation and treatment outcomes across etiologies in a 3-month consecutive keratitis cohort.**

**Abbreviations:** CI, confidence interval; OR, odds ratio.

**Footnotes:** *a* **Presentation** indicates disease severity at the time of initial clinical evaluation based on ulcer size and depth, classified into ‘non-severe’ or ‘severe’ and compared between keratitis types by Fisher’s exact test, *P* value is shown. *b* **Clinical outcome** indicates treatment response as ‘healed’ (infection resolution with corneal healing) or ‘unhealed’ (non-response to medical therapy requiring TPK) compared between keratitis types by Fisher’s exact test, *P* value is shown. *c* **Ref.** indicates the reference category used for odds ratio calculations for presentation and clinical outcome among keratitis of different etiologies *d* **Odds ratios for presentation** represent the likelihood of ‘severe’ presentation, calculated using a univariate unadjusted model. *e* **Odds ratios for clinical outcome** represent the likelihood of ‘unhealed’ outcome, calculated using a univariate unadjusted model. Significance defined as *P* < .05.

**Supplementary Table 5.**

| **Keratitis** | **Outcome^a^** | **Healed- Vision^b^**  **(*P*<0.001)** | | **OR (95% CI)^d^** | ***P* Value** | **OR (95% CI)^d^** | ***P* Value** |
| --- | --- | --- | --- | --- | --- | --- | --- |
| *A. flavus* (n=420) | Healed (59.52%) | Good vision (54.0%) | Poor Vision (46.0%) | 3.52 (2.47-5.03) | <0.001 | 1.68 (1.20-2.33) | 0.002 |
| *Fusarium* spp*.* (n=481) | Healed (77.96%) | Good Vision (80.53%) | Poor Vision (19.47%) | Ref. ^c^ | - | 0.48 (0.34-0.67) | <0.001 |
| Culture -ve (n=403) | Healed (90.57%) | Good Vision (66.30%) | Poor Vision (33.70%) | 2.10 (1.50-2.94) | <0.001 | Ref. ^c^ | - |

**Supplementary Table 5. Comparison of treatment and visual outcomes across etiologies of fungal keratitis.**

**Abbreviations:** CI, confidence interval; OR, odds ratio; TPK, therapeutic penetrating keratoplasty.

**Footnotes:**
^a^ **Outcome** indicates response to medical therapy, as ‘healed’ (resolution of infection with corneal healing). ^b^ **Visual outcome of healed eyes** indicates visual outcome among patients who achieved corneal healing, classified as ‘good vision’ or ‘poor vision’. Visual outcomes were compared across keratitis etiologies using the χ² test, and the overall *P* value is shown. ^c^ **Ref.** indicates the reference category used for odds ratio calculations. *Fusarium* spp. and culture-negative keratitis were used as reference categories in separate analyses. ^d^ **Odds ratios** represent the likelihood of poor vision among patients with healed keratitis, calculated using a univariate unadjusted model, with the indicated reference category. Significance was defined as *P* < .05**.**

**Supplementary Table 6.**

|  | **Ulcer Position** | **Central (%)** | **Paracentral (%)** | **Peripheral (%)** | ***P* value^a^** |
| --- | --- | --- | --- | --- | --- |
| **Total-Healed** | *A. flavus* (n=30) | 30.00 | 46.67 | 23.33 | 0.994 |
|  | *Fusarium* sp. (n=30) | 33.33 | 46.67 | 20.00 |  |
|  | Culture -ve (n=30) | 33.33 | 43.33 | 23.33 |  |
| **Healed- Good Vision** | *A. flavus* (n=19) | 47.37 | 36.84 | 15.79 | 0.522 |
|  | *Fusarium sp*. (n=18) | 22.22 | 50.00 | 27.78 |  |
|  | Culture -ve (n=18) | 27.78 | 55.56 | 16.67 |  |
| **Healed- Poor vision** | *A. flavus* (n=11) | 0 | 63.64 | 36.36 | 0.032 |
|  | *Fusarium* sp. (n=12) | 50.00 | 41.67 | 8.33 |  |
|  | Culture -ve (n=12) | 41.67 | 25.00 | 33.33 |  |

**Supplementary Table 6.** **Comparison of ulcer location across etiologies among healed patients.**

**Footnotes:** a ***P* value** determined using the Fisher’s exact test. Significance defined as *P* < .05.

**Supplementary Table 7.**

| **Gene ID** | **Gene Name** | **log2 Fold Change**  **(-log10 *P* value)** | | | **Gene function** |
| --- | --- | --- | --- | --- | --- |
|  |  | **Ctrl**  **Vs**  **NS-H** | **NS-H**  **Vs**  **NS-UH** | **NS-H**  **vs**  **S-UH** |  |
| *IL-17C* | Interleukin-17C | 7.13  (4.84) | -4.38  (3.02) | -4.99  (1.67) | Pro-inflammatory cytokine involved in epithelial innate immune responses and antimicrobial defense. |
| *FAM53A* | Family with sequence similarity 53, member A | 6.10  (3.75) | -3.73  (2.70) | -4.93  (1.67) | Nuclear protein associated with *Wnt* signaling and regulation of cell proliferation. |
| *LCE3E* | Late Cornified Envelope 3E | 6.11  (3.52) | -3.74  (2.40) | -4.93  (1.58) | Epidermal structural protein involved in keratinocyte differentiation and skin barrier formation. |
| *SLC26A4* | Solute Carrier Family 26 Member 4 | -7.85  (5.58) | 6.86  (3.82) | 8.90  (5.33) | Anion transporter (pendrin) regulating iodide, chloride, and bicarbonate transport in epithelial tissues. |
| *EXO1* | Exonuclease 1 | -5.53  (3.39) | 5.70  (3.14) | 6.49  (3.34) | DNA exonuclease involved in mismatch repair, recombination, and maintenance of genome stability. |
| *ROBO4* | Roundabout Homolog 4 | -5.61  (3.36) | 5.47  (2.86) | 5.62  (2.39) | Endothelial receptor regulating angiogenesis and vascular integrity. |
| *LILRB4* | Leukocyte Immunoglobulin-Like Receptor B4 | -3.39  (1.31) | 6.16  (3.90) | 6.08  (3.12) | Inhibitory immune receptor on myeloid cells that suppresses immune activation. |

**Supplementary Table 7. Candidate healing-associated signatures in *Fusarium* keratitis**.

Fold change (log2) of gene expression and significance (-log10 *P* value) are shown for respective comparisons, along with gene function.

**Abbreviations:** Ctrl, Healthy control (cadaveric cornea); NS-H, Non-severe healed; NS-UH, Non-severe unhealed; S-UH, Severe unhealed groups.

**Supplementary Table 8.**

| **Gene ID** | **Gene Name** | **log2 Fold Change**  **(-log10 *P* value)** | | | | **Gene Function** |
| --- | --- | --- | --- | --- | --- | --- |
|  |  | **Ctrl**  **vs**  **NS-UH** | **Ctrl**  **vs**  **S-UH** | **NS-H**  **Vs**  **NS-UH** | **NS-H**  **vs**  **S-UH** |  |
| *FOLR3* | Folate Receptor Gamma | 6.89  (5.01) | 7.14  (4.52) | 9.09  (3.76) | 6.35  (3.37) | Secreted folate-binding protein involved in folate transport and immune modulation. |
| *LRAT* | Lecithinretinol acyltransferase | 3.36  (2.18) | 3.40  (1.56) | 5.00  (2.67) | 5.04  (2.08) | Enzyme that converts retinol to retinyl esters for vitamin A storage. |
| *MEDAG* | Mesenteric Estrogen-Dependent Adipogenesis | 4.83  (3.05) | 6.92  (5.42) | 5.00  (3.01) | 7.08  (5.14) | Regulates adipocyte differentiation and lipid accumulation. |
| *PNPLA3* | Patatin-like phospholipase domain-containing protein 3 | 3.50  (4.85) | 3.83  (4.29) | 3.77  (3.60) | 4.10  (3.31) | Lipid metabolism enzyme involved in triglyceride remodeling and hydrolysis. |
| *CES1* | Carboxylesterase 1 | -3.67  (4.23) | -5.88  (2.64) | -2.77  (2.08) | -4.98  (1.76) | Enzyme that hydrolyzes esterified lipids, xenobiotics, and drugs. |
| *ICOSL* | Inducible T-cell Co-Stimulator Ligand | -8.65  (8.03) | -6.41  (3.09) | -9.35  (8.22) | -7.11  (3.29) | Immune co-stimulatory ligand that activates ICOS-expressing T cells. |
| *CREB5^#^* | cAMP responsive element binding protein 5 | - | 2.48  (2.94) | 3.69  (3.83) | 4.90  (8.82) | Transcription factor regulating gene expression in response to cAMP signaling. |
| *PDE4B^#^* | Phosphodiesterase-4B | - | 2.49  (5.07) | 2.56  (3.57) | 5.58  (19.11) | Hydrolyzes cAMP to regulate inflammatory and immune signaling pathways. |

**Supplementary Table 8. Candidate unhealing-associated signatures in *Fusarium* keratitis**.

Fold change (log2) of gene expression and significance (-log10 *P* value) are shown for respective comparisons, along with gene function. ‘#’ indicates genes selected from Fu_983 subset (Supplementary Figure 2C).

**Abbreviations:** Ctrl, Healthy control (cadaveric cornea); NS-H, Non-severe healed; NS-UH, Non-severe unhealed; S-UH, Severe unhealed groups.

**Supplementary Table 9.**

| **Gene ID** | **Gene Name** | **log2 Fold Change**  **(-log10 *P* value)** | | | **Gene functions** |
| --- | --- | --- | --- | --- | --- |
|  |  | **Ctrl vs NS-H** | **Ctrl**  **vs**  **S-UH** | **NS-H vs**  **S-UH** |  |
| *C5AR1* | Complement Component 5a Receptor 1 | -1.43  (1.68) | 4.32  (17.97) | 5.75  (29.43) | G protein–coupled receptor mediating inflammatory responses to complement C5a. |
| *RGS2* | Regulator of G protein signaling 2 | -3.48  (5.04) | 2.77  (4.03) | 6.25  (17.29) | Modulates GPCR signaling by accelerating GTPase activity of Gα proteins. |
| *PDE4B* | Phosphodiesterase-4B | -3.09  (5.37) | 2.49  (5.07) | 5.58  (19.11) | Hydrolyzes cAMP to regulate inflammatory and immune signaling pathways. |
| *LY96* | Lymphocyte antigen 96 | -2.76  (3.68) | 2.54  (5.38) | 5.30  (15.12) | Co-receptor for *TLR4* in innate immunity. |
| *LRRK2* | Leucine-rich repeat kinase 2 | -2.11  (2.71) | 3.01  (7.41) | 5.12  (17.25) | Multifunctional kinase involved in vesicle trafficking, immune signaling, and neuronal function. |
| *CCRL2* | C-C motif chemokine receptor-like 2 | -2.52  (3.40) | 1.71  (2.41) | 4.23  (10.95) | Atypical chemokine receptor involved in leukocyte migration and inflammatory regulation. |
| *IL2RG* | Interleukin 2 Receptor Subunit Gamma | -1.20  (1.46) | 3.32  (13.81) | 4.51  (22.46) | Common γ-chain shared by multiple cytokine receptors essential for lymphocyte development. |
| *IFITM2* | Interferon-Induced Transmembrane Protein 2 | -2.24  (3.99) | 3.04  (8.81) | 5.28  (20.08) | Antiviral membrane protein that restricts viral entry into host cells. |
| *MYO1F* | Myosin1F | -2.13  (3.12) | 3.55  (11.28) | 5.67  (24.48) | Actin-based motor protein involved in immune cell adhesion, migration, and phagocytosis. |
| *SLC16A3* | Solute Carrier Family 16 Member 3 | 1.06  (2.96) | 2.64  (20.81) | 1.58  (7.03) | Proton-linked monocarboxylate transporter exporting lactate from glycolytic cells. |
| *ITGAL* | Integrin subunit alpha L | -2.35  (2.26) | 2.64  (4.74) | 4.99  (10.99) | Leukocyte adhesion receptor forming *LFA-1* complex for immune cell adhesion and migration. |
| *SRGN* | Serglycin | -1.18  (1.34) | 4.40  (19.53) | 5.59  (28.79) | Proteoglycan involved in storage and secretion of inflammatory mediators in immune cells. |

**Supplementary Table 9. Candidate severity/outcome-associated signatures in *A. flavus* keratitis**.

Fold change (log2) of gene expression and significance (-log 10 *P* value) are shown for respective comparisons, along with gene function.

**Abbreviations:** Ctrl, Healthy control (cadaveric cornea); NS-H, Non-severe healed; NS-UH, Non-severe unhealed; S-UH, Severe unhealed groups.

**Supplementary Table 10.**

| **Characteristic** | **Total (n=102)** | ***A . flavus* Keratitis**  **(n= 24)** | ***Fusarium* keratitis**  **(n=54)** | **Culture -ve keratitis**  **(n=10)** | **Others**  **(n=14)** | ***P* value** |
| --- | --- | --- | --- | --- | --- | --- |
| Male participants, No. (%) | 64 (62.7) | 18 (75.0) | 34 (63.0) | 4 (40.0) | 8 (57.1) | 0.241^$^ |
| Female participants, No. (%) | 38 (37.3) | 6 (25.0) | 20 (37.0) | 6 (60.0) | 6 (42.9) |  |
| Age, mean (SD), y | 50.25 (16.3) | 55.67 (18.6) | 47.74 (14.4) | 47.30 (15.1) | 52.92 (18.6) | 0.201^#^ |
| Severe, No. (%) | 37 (36.3) | 14 (58.3) | 19 (35.2) | 1 (10.00) | 3 (21.4) | 0.016^$^ |
| Healed, No. (%) | 68 (67.7) | 10 (41.7) | 38 (70.37) | 10 (100.0) | 11 (78.5) | 0.024^$^ |
| Unhealed, No. (%) | 22 (21.6) | 11 (45.8) | 9 (16.7) | 0 | 2 (14.3) |  |
| LOF, No. (%) | 11 (10.7) | 3 (12.5) | 7 (13.0) | 0 | 1 (7.12) |  |
| Age range, y | 13 – 85 | 17 – 85 | 13 – 75 | 27 – 66 | 29 – 75 | - |
| Diabetes, No. (%) | 22 (21.6) | 8 (33.3) | 6 (11.1) | 3 (30.0) | 5 (35.7) | 0.027^$^ |
| Hypertension, No. (%) | 14 (13.7) | 5 (20.8) | 6 (11.1) | 2 (20.0) | 1 (7.2) | 0.572^$^ |
| Cardiac, No. (%) | 5 (4.9) | 3 (12.5) | 2 (3.7) | 0 | 0 | 0.301^$^ |

**Supplementary Table 10. Demographic and clinical characteristics of the validation study cohort.**

Baseline demographics, systemic comorbidities, clinical characteristics and treatment outcomes for 102 patients in the validation cohort, stratified by infecting organism (*A flavus*, *Fusarium* spp., no growth, and others). Categorical data are presented as No. (%) and continuous variables as mean (SD). *P* values were determined using the Fisher’s exact test (indicated by $) or 1-way ANOVA (indicated by #). Significance was defined as *P* < .05. LOF indicates loss of follow-up.

**Supplementary Table 11.**

| **Gene** | **AUC** | ***P* value** | **Cut off fold change** | **Sensitivity % 95% CI** | **Specificity % 95% CI** |
| --- | --- | --- | --- | --- | --- |
| *PDE4B* | 0.8846 | 0.0019 | >29.64 | 92.31% (66.69% to 99.61%) | 70.00% (39.68% to 89.22%) |
| *LY96* | 0.9077 | 0.0010 | > 9.588 | 84.62% (57.77% to 97.27%) | 90.00% (59.58% to 99.49%) |
| *LRRK2* | 0.9385 | 0.0004 | > 9.830 | 84.62% (57.77% to 97.27%) | 100.0% (72.25% to 100.0%) |
| *SRGN* | 0.8769 | 0.0024 | > 58.82 | 76.92% (49.74% to 91.82%) | 80.00% (49.02% to 96.45%) |
| *RGS2* | 0.8231 | 0.0092 | > 5.567 | 76.92% (49.74% to 91.82%) | 80.00% (49.02% to 96.45%) |
| *CCRL2* | 0.8769 | 0.0024 | >25.64 | 76.92% (49.74% to 91.82%) | 100.00% (72.25% to 100.00%) |
| *C5AR1* | 0.9615 | 0.0002 | >32.66 | 84.62% (57.77% to 97.27%) | 100.00% (72.25% to 100.0%) |
| *MYO1F* | 0.9385 | 0.0004 | >43.20 | 76.92% (49.64% to 91.82%) | 100.0% (72.25% to 100.0%) |

**Supplementary Table 11. Area under receiver operating characteristic analysis of severity-associated signatures.**

AUROC analysis of fold changes in normalized gene expression for a validation cohort of 23 patients with fungal keratitis (*Fusarium* keratitis, n = 9; *A. flavus* keratitis, n = 14). The analysis evaluates the diagnostic performance of candidate gene signatures to differentiate severe from non-severe cases. For each signature, the area under the curve (AUC), *P* values, optimal cut-off fold changes, and corresponding sensitivity and specificity with 95% CIs are shown.

**Supplementary Table 12.**

| **Patient ID** | **Pathogen** | **Ulcer Size (cm^2^)** | **Depth**  **(%)** | **Initial Visit** | | | **Final Visit** | | ***LY96*** | ***LRRK2*** | ***SRGN*** | ***RGS2*** | ***PDE4B*** | ***CCRL2*** | ***C5AR1*** | ***MYO1F*** | **Outcome cluster** |
| --- | --- | --- | --- | --- | --- | --- | --- | --- | --- | --- | --- | --- | --- | --- | --- | --- | --- |
|  |  |  |  | **Presentation** | **UCVA** | | **Outcome** | **UCVA** |  |  |  |  |  |  |  |  |  |
| **A8** | ***A. Flavus*** | **20** | **50** | **severe** | **FCF** | **TPK^+++^** | | **HM** | **104.68** | **252.69** | **1,019.59** | **168.44** | **1,207.79** | **281.70** | **749.73** | **866.27** | Poor  (Inflammation high) |
| **A3** | ***A. Flavus*** | **25** | **NA** | **severe** | **FCF** | **TPK^++^** | | **HM** | **358.34** | **586.72** | **1,774.66** | **592.76** | **722.19** | **77.14** | **425.41** | **1,523.49** |  |
| A4 | *A. Flavus* | 49 | 75 | severe | HM | TPK | | FCF | 209.44 | 141.17 | 661.06 | 217.00 | 221.65 | 62.63 | 148.33 | 173.03 |  |
| A5 | *A. Flavus* | 30 | 85 | severe | FCF | TPK^+^ | | HM | 118.25 | 160.83 | 570.40 | 239.24 | 341.60 | 16.08 | 543.60 | 341.94 |  |
| A6 | *A. Flavus* | 49 | 80 | severe | HM | TPK^+^ | | LP+ | 132.13 | 50.02 | 544.54 | 77.90 | 190.18 | 169.98 | 144.16 | 112.73 |  |
| A1 | *A. Flavus* | 49 | 80 | severe | HM | TPK^++^ | | LP+ | 230.35 | 11.84 | 455.52 | 82.84 | 141.53 | 141.89 | 9.72 | 13.04 |  |
| A14 | *A. Flavus* | 25 | 50 | severe | NLP | TPK^+^ | | NLP | 34.94 | 30.65 | 557.98 | 8.97 | 610.46 | 295.05 | 101.33 | 151.44 |  |
| A9 | *A. Flavus* | 18 | 50 | severe | HM | LOF-UH | | HM | 18.46 | 12.75 | 226.22 | 6.84 | 222.18 | 154.37 | 301.66 | 72.42 |  |
| A2 | *A. Flavus* | 20 | 40 | severe | HM | TPK^++^ | | HM | 36.32 | 6.54 | 100.56 | 20.35 | 46.05 | 21.48 | 7.62 | 8.97 |  |
| U2 | *A. Flavus* | 30 | 40 | severe | LP+ | TPK | | LP+ | 18.39 | 26.21 | 151.72 | 13.73 | 19.81 | 9.95 | 39.16 | 157.38 |  |
| A11 | *A. Flavus* | 6 | 40 | Non-severe | 6/36 | Healed | | 5/60 | NA | 58.26 | 80.10 | 14.13 | 92.74 | NA | 46.49 | 146.39 | Better (Inflammation low) |
| A13 | *A. Flavus* | 20.25 | 50 | severe | 6/60 | Healed | | 6/36 | 43.46 | 43.58 | 59.72 | 5.70 | 43.28 | 29.81 | 63.99 | 79.84 |  |
| A7 | *A. Flavus* | 7 | 20 | Non-severe | 6/9 | LOF-H | | 6/9 | 5.67 | 7.82 | 57.91 | 9.58 | 44.07 | 13.70 | 31.05 | 23.88 |  |
| A12 | *A. Flavus* | 36 | 50 | severe | 5/60 | Healed | | 6/36 | NA | NA | 15.58 | 3.52 | 6.75 | NA | 11.66 | 15.16 |  |
| A15 | *A. Flavus* | 9 | 30 | Non-severe | 6-9P | Healed | | 6/9P | 9.08 | 2.95 | 9.56 | 1.84 | 3.75 | 2.76 | 3.90 | 6.46 |  |
| A10 | *A. Flavus* | 7.29 | NA | Non-severe | HM | LOF-H | | NA | 3.39 | 3.98 | 18.08 | 5.44 | 24.89 | 19.32 | 11.75 | 15.86 |  |

**Supplementary Table 12. Gene expression profiles demonstrating outcome-associated clustering patterns in A. flavus keratitis.**

This table summarizes clinical features, ulcer characteristics, and relative gene expression (fold change over healthy cadaveric controls) of corneal-response markers *(LY96, LRRK2, SRGN, RGS2, PDE4B, CCRL2, C5AR1, MYO1F*) in a subset of patients with Aspergillus flavus keratitis at presentation for whom expression data were available for majority of genes. Visual acuity (UCVA) at initial and final visits and clinical outcomes, including requirement for therapeutic penetrating keratoplasty (TPK), are presented. Patients are categorized based on clinical outcome: healed, and TPK subgroups (TPK+, perforation; TPK++, graft failure or reinfection; TPK+++, evisceration or phthisis). LOF indicates loss of follow-up at Aravind Eye Hospital, Madurai, with outcomes confirmed via telephonic inquiry. Predefined thresholds for high gene expression were applied (*LY96* > 9.588; *LRRK2* > 9.83; *SRGN* > 58.82; *RGS2* > 5.567; *PDE4B* > 29.64; *CCRL2* >25.64; *C5AR1* > 32.66; *MYO1F* > 43.20), and color-coding was used to indicate elevated expression levels (Pink: > cutoff; Red: >5x cutoff). Gene expression analysis was performed on microbiologically confirmed Aspergillus flavus cases, with clinical data obtained subsequently. Patients requiring TPK and post TPK complications demonstrate markedly elevated expression across multiple inflammatory and immune activation genes. In contrast, patients with healed outcomes generally exhibit lower overall gene-expression levels. Descending order of visual acuity: 6/9 → 6/9P → 6/36 → 6/60 → 5/60 → FCF → HM → LP+ → NLP.

**Abbreviations:** HM, Hand Movements; FCF, Counting Fingers; LOF, Loss of Follow-up; LOF-H/UH, Loss of Follow-up- Healed/Unhealed; HM, Hand Movements; NLP, No Light Perception; LP+, Light Perception; NA, Data not available; (UCVA)P, Uncorrected visual acuity partial ; TPK, Therapeutic Penetrating Keratoplasty.

**Supplementary Table 13.**

| **Patient ID** | **Pathogen** | **Ulcer Size (mm^2^)** | **Depth**  **(%)** | **Initial Visit** | | **Final Visit** | | ***ICOSL*** | ***SLC26A4*** | ***LY96*** | ***LRRK2*** | ***SRGN*** | ***RGS2*** | ***PDE4B*** | **Outcome cluster** |
| --- | --- | --- | --- | --- | --- | --- | --- | --- | --- | --- | --- | --- | --- | --- | --- |
|  |  |  |  | **Presentation** | **UCVA** | **Outcome** | **UCVA** |  |  |  |  |  |  |  |  |
| F2 | *Fusarium* | 16 | 30 | Severe | 6/18P | Healed | 6/6 | 2.99 | 1.48 | 1.72 | 0.34 | 9.35 | 0.53 | 12.57 | Good |
| F5 | *Fusarium* | 3 | NA | Non-severe | 3/60 | Healed | 6/18 | 4.36 | 1.88 | 0.89 | 0.91 | 2.96 | 0.39 | 12.15 |  |
| F6 | *Fusarium* | 17.5 | 60 | Severe | HM | Healed | 3/60 | 3.99 | 3.01 | 5.17 | 2.73 | 41.81 | 0.89 | 34.39 |  |
| F7 | *Fusarium* | 2 | 30 | Non-severe | 6/6 | Healed | 6/6 | 8.10 | 0.30 | 10.09 | 15.83 | 7.56 | 0.72 | 35.85 |  |
| F8 | *Fusarium* | 2 | 20 | Non-severe | 6/24P | Healed | 6/6P | 7.17 | 1.17 | 1.01 | 1.95 | 1.32 | 0.26 | 6.24 |  |
| F9 | *Fusarium* | 9.63 | 30 | Non-severe | 6/60 | Healed | 6/24 | 3.51 | 4.16 | 4.21 | 2.55 | 33.57 | 2.44 | 21.64 |  |
| A17 | *A. Flavus* | 2.25 | NA | severe | 5/60 | Healed | 6/36 | 2.89 | 5.95 | 10.39 | 0.04 | 24.57 | 0.67 | 9.00 |  |
| F4 | *Fusarium* | 7.5 | 55 | Non-severe | 3/60 | Healed | 3/60 | 1.81 | 0.94 | 1.73 | 1.01 | 3.88 | 0.41 | 3.41 | Variable |
| F3 | *Fusarium* | 29.12 | 20 | Severe | 6/36 | Healed | 6/18 | 0.74 | 3.22 | 7.34 | 5.13 | 62.51 | 4.02 | 102.10 |  |
| U1 | *Fusarium* | 30 | NA | Severe | 1/60 | TPK | HM | 2.27 | 15.62 | 4.51 | 4.92 | 23.89 | 3.94 | 53.21 |  |
| A18 | *A. Flavus* | 4 | 60 | Non-severe | ½/60 | LOF | ½/60 | 1.40 | 0.73 | 65.24 | NA | 17.05 | 0.51 | 6.90 |  |
| U2 | *A. Flavus* | 30 | 40 | severe | LP+ | TPK | LP+ | 0.41 | 0.43 | 18.39 | 26.21 | 151.72 | 13.73 | 19.81 | Poor |
| A19 | *A. Flavus* | 22.5 | NA | Severe | 5/60 | TPK | HM | 8.17 | 19.53 | 46.39 | 36.80 | 566.74 | 25.24 | 1,094.56 |  |

**Supplementary Table 13. Gene expression profiles demonstrating outcome-associated clustering patterns in *Fusarium* and A. flavus keratitis.**

This table summarizes clinical features, ulcer characteristics, and relative gene expression (fold change over healthy cadaveric controls) of *ICOSL, SLC26A4, LY96, LRRK2, SRGN, RGS2*, and *PDE4B* in a subset of patients with *Fusarium* or Aspergillus flavus keratitis at presentation for whom expression data were available. Visual acuity (UCVA) at initial and final visits and clinical outcomes, including requirement for therapeutic penetrating keratoplasty (TPK), are presented. LOF indicates loss to follow-up at Aravind Eye Hospital, Madurai. Predefined thresholds for high gene expression were applied (*ICOSL* > 2.7; *SLC26A4* > 2.6; *LY96* > 9.588; *LRRK2* > 9.83; *SRGN* > 58.82; *RGS2* > 5.567; *PDE4B* > 29.64), with color-coding used to indicate elevated expression (pro-inflammatory genes: pink, >cutoff; red, >5× cut-off), ICOSL upregulation (green, > cut-off), and SLC26A4 expression (orange, > cutoff). Gene expression analysis was performed on microbiologically confirmed Fusarium and Aspergillus flavus cases, with clinical data obtained subsequently. Patients with good outcomes show relatively higher ICOSL expression with low inflammatory markers, whereas poor outcomes are characterized by elevated inflammatory gene expression; variable outcomes show low to intermediate expression profiles. Elevated SLC26A4 expression is noted primarily in patients with poorer initial visual acuity. Descending order of visual acuity: 6/6 → 6/6P → 6/9 → 6/9P → 6/18 →6/18P → 6/24 → 6/24P→ 6/36 → 6/60 → 5/60 →3/60 → ½/60 → FCF → HM → LP+

**Abbreviations:** HM, Hand Movements; LOF, Loss of Follow-up; HM, Hand Movements; LP+, Light Perception; NA, Data not available; (UCVA)P, Uncorrected visual acuity partial ; TPK, Therapeutic Penetrating Keratoplasty.

**Supplementary Table 14.**

| **Patient ID** | **Pathogen** | **Ulcer Size (cm^2^)** | **Depth**  **(%)** | **Initial Visit** | | **Final Visit** | | ***ICOSL*** | ***LY96*** | ***LRRK2*** | ***SRGN*** | ***RGS2*** | ***PDE4B*** | **Outcome Cluster** |
| --- | --- | --- | --- | --- | --- | --- | --- | --- | --- | --- | --- | --- | --- | --- |
|  |  |  |  | **Presentation** | **UCVA** | **Outcome** | **UCVA** |  |  |  |  |  |  |  |
| C1 | *Curvularia* | 14 | NA | Non-severe | 6/12 | Healed | 6/12P | 10.42 | 0.77 | 2.20 | 15.37 | 0.60 | 44.89 | **Good**  **(Clinical & visual)** |
| C2 | *Curvularia* | 4 | 30 | Non-severe | 6/6 | Healed | 6/6 | 32.88 | 0.73 | 0.24 | 1.61 | 0.12 | 3.65 |  |
| C4 | *Curvularia* | 6.25 | NA | Non-severe | 6/18 | Healed | 6/12 | 7. 19 | 0.97 | 0.74 | 8.34 | 0.10 | 62.99 |  |
| N5 | Culture -ve | 25 | 20 | Non-severe | 6/36 | Healed | 6/12P | 10.09 | 0.56 | 7.35 | 1.32 | 0.08 | 32.82 |  |
| N8 | Culture -ve | 9 | NA | Non-severe | 6/18P | Healed | 6/12P | 17.97 | 1.15 | 2.86 | 3.04 | 0.39 | 4.87 |  |
| C3 | *Curvularia* | 2.25 | 60 | Non-severe | 5/60 | Healed | 6/60 | 6.38 | 1.57 | 1.73 | 2.71 | 0.12 | 4.09 |  |
| P4 | *Pseudomonas* | 15 | 70 | Severe | FCF | Healed | 6/24 | 6.99 | 4.44 | 16.60 | 29.16 | 2.28 | 17.53 |  |
| N7 | Culture -ve | 10.5 | 30 | Non-severe | 6/12 | Healed | 6/9 | 6.97 | 11.07 | 27.73 | 66.39 | 6.44 | 66.96 |  |
| C6 | *Curvularia* | 1.2 | NA | Non-severe | 6/9 | Healed | 6/6P | 0.52 | 1.12 | 0.97 | 4.98 | 0.40 | 6.81 | **variable (visual)** |
| C7 | *Curvularia* | 1.5 | NA | Non-severe | 6/9 | Healed | 6/6P | 0.90 | 2.36 | 2.78 | 20.59 | 0.29 | 20.58 |  |
| P1 | *Pseudomonas* | 4 | 30 | Non-severe | 6/18P | Healed | 6/18P | 0.26 | 0.73 | 0.17 | 3.34 | 0.34 | 3.66 |  |
| P2 | *Pseudomonas* | 4 | NA | Non-severe | 6/60 | Healed | 5/60 | 1.84 | 4.37 | 0.14 | 1.67 | 0.04 | 1.82 |  |
| P6 | *Pseudomonas* | 12 | 30 | Non-severe | 6/60 | LOF | ½/60 | 2.35 | 2.61 | 0.76 | 3.05 | 0.32 | 1.50 |  |
| P3 | *Pseudomonas* | 4 | 30 | Non-severe | 6/6 | Healed | 6/12P | 2.13 | 1.73 | 2.80 | 24.12 | 1.16 | 38.75 |  |
| N2 | Culture –ve | 2.5 | NA | Non-severe | 5/60 | Healed | 5/60 | 0.11 | 24.58 | 4.50 | 6.52 | 0.84 | 1.78 |  |
| N4 | Culture -ve | 9 | 90 | severe | 6/60 | Healed | 6/18 | 1.16 | 36.71 | 28.49 | 235.06 | 20.10 | 133.00 |  |
| N3 | Culture -ve | 1.25 | NA | Non-severe | 6/6 | Healed | 6/6 | 2.42 | 35.25 | 80.03 | 133.96 | 17.88 | 42.70 |  |
| N1 | Culture –ve | 25 | NA | severe | FCF | Healed | ½/60 | 0.67 | 72.46 | 151.15 | 137.64 | 30.86 | 80.20 | **Poor (visual)** |
| N6 | Culture -ve | 1 | NA | Non-severe | 1/60 | Healed | ½/60 | 6.75 | 31.06 | 96.35 | 211.10 | 39.07 | 129.16 |  |
| **C5*** | ***Curvularia*** | **30** | **50** | **Non-severe** | **1/2/60** | **Unhealed** | **Evis.** | **0.73** | **329.96** | **517.21** | **2175.95** | **133.06** | **2463.91** | **Worst** |
| **P5*** | ***Pseudomonas*** | **64** | **80** | **Severe** | **LP+** | **Unhealed** | **LP+** | **2.81** | **62.25** | **147.19** | **1025.47** | **64.32** | **889.52** |  |

**Supplementary Table 14. Gene expression profiles demonstrating outcome-associated clustering patterns in microbial keratitis.**

This table summarizes clinical features, microbiological etiology, and relative gene expression (fold change over healthy cadaveric controls) of *ICOSL*, *LY96, LRRK2, SRGN, RGS2*, and *PDE4B* in a subset of patients with microbial keratitis at presentation for whom expression data were available for majority of the listed genes. Visual acuity (UCVA) at initial and final visits and clinical outcomes are presented. LOF indicates loss to follow-up at Aravind Eye Hospital, Madurai. Predefined thresholds for high gene expression were applied (*ICOSL* > 2.7; *LY96* > 9.588; *LRRK2* > 9.83; *SRGN* > 58.82; *RGS2* > 5.567; *PDE4B* > 29.64), with color-coding to indicate elevated expression (pro-inflammatory genes; Pink: >cutoff, Red: >5× cutoff) and ICOSL upregulation (Green: > cutoff). Gene expression analysis was performed on microbiologically confirmed cases, with clinical data obtained subsequently. Patients in the poor and worst outcome clusters demonstrate elevated expression of multiple inflammatory genes (*LY96, LRRK2, SRGN, PDE4B*). Patients with good clinical and visual outcomes primarily exhibit relatively higher *ICOSL* expression with low inflammatory markers. The **variable outcome group** includes patients with low to intermediate gene expression profiles. Descending order of visual acuity: 6/6 → 6/6P → 6/9 → 6/9P → 6/12 → 6/12P → 6/18 → 6/18P → 6/24 → 6/36 → 6/60 → 5/60 → 1/60 → 0.5/60 → FCF → LP+

**Abbreviations:** LOF, Loss of Follow-up; LP+, Light Perception; NA, Data not available; (UCVA)P, Uncorrected visual acuity, partial; TPK, Therapeutic Penetrating Keratoplasty; Evis., evisceration.

**Supplementary Table 15.**

| **Patient ID** | ***ICOSL*** | ***SLC26A4*** | ***LY96*** | ***LRRK2*** | ***SRGN*** | ***RGS2*** | ***PDE4B*** | ***CCRL2*** | ***C5AR1*** | ***MYO1F*** | **Microbiology Report** |
| --- | --- | --- | --- | --- | --- | --- | --- | --- | --- | --- | --- |
| U1 | 2.3 | 15.6 | 4.5 | 4.9 | 23.9 | 3.9 | 60.3 | 11.4 | 12.0 | 25.6 | *Fusarium spp*. |
| U2 | 0.4 | 0.4 | 18.4 | 26.2 | 151.7 | 13.7 | 19.8 | 10.0 | 39.2 | 157.4 | *A. flavus* |
| U3 | 5.8 | 5.4 | 12.2 | 6.5 | 14.0 | 2.2 | 4.7 | 12.9 | 6.3 | 15.7 | Culture -ve |
| U4 | 11.8 | 32.0 | 30.1 | 49.6 | 384.4 | 18.8 | 882.5 | 228.7 | 216.3 | 461.0 | *Fusarium spp.* |
| U5 | 4.2 | 61.1 | 74.8 | 65.3 | 532.7 | 19.2 | 348.5 | 59.0 | 126.4 | 391.3 | Culture -ve |
| U6 | 1.3 | 33.6 | 47.8 | 36.1 | 256.7 | 22.8 | 168.4 | 60.6 | 28.8 | 94.9 | *P. aeruginosa*+ unknown fungus |

**Supplementary Table 15. Presentation-stage host-response gene expression profiles in six masked keratitis samples.**

Relative expression (fold change) of ICOSL, SLC26A4, LY96, LRRK2, SRGN, RGS2, PDE4B, CCRL2, C5AR1, and MYO1F was analyzed while masked to microbiological diagnosis and clinical characteristics. Predefined gene-specific cutoffs were applied (ICOSL >2.7; SLC26A4 >2.6; LY96 >9.588; LRRK2 >9.83; SRGN >58.82; RGS2 >5.567; PDE4B >29.64; CCRL2 >25.64; C5AR1 >32.66; MYO1F >43.20), with color coding indicating expression relative to these thresholds. Microbiological diagnoses, revealed after expression analysis, are shown for reference.

**Supplementary Table 16.**

| **Etiology** | ***ICOSL*** | ***SLC26A4*** | **8-gene severity profile** | **Masked outcome interpretation** |
| --- | --- | --- | --- | --- |
| ***Fusarium* spp.** | Low | High or low | Moderate - high | Unhealed |
|  | High | High or low | Low-moderate | Healed |
|  | High | High | Moderate - high | Unfavorable course; variable/indefinite |
| ***A. flavus*** | High or Low | High or low | Moderate - very high | Unhealed |
|  | Low | High or low | Low-moderate | Unfavorable course; Variable/indefinite |
|  | High | High or low | Low | Healed |
| **Culture-negative** | High | High or low | Low - high | Healed |
|  | Low | High or low | Low - High | Unfavorable course; Healed |
|  | Low | High or low | Very high | Unfavorable course; Variable/indefinite |

**Supplementary Table 16. Candidate framework for interpretation of presentation-stage host-response profiles**

Candidate criteria were derived from observed patient-level expression patterns to provide a provisional framework for etiology-specific outcome interpretation and require validation in larger cohorts. High and low indicate expression above and below predefined gene-specific cutoffs, respectively. Cutoffs were ICOSL >2.7 and SLC26A4 >2.6; cutoffs for the 8-gene severity-associated panel were LY96 >9.588, LRRK2 >9.83, SRGN >58.82, RGS2 >5.567, PDE4B >29.64, CCRL2 >25.64, C5AR1 >32.66, and MYO1F >43.20. Low, moderate, high, and very high denote the combined number and magnitude of upregulation of the 8 severity-associated genes above their predefined cutoffs. Outcome interpretation was based on the combined ICOSL, SLC26A4, and 8-gene severity profile within each etiology. **Unhealed**, requiring therapeutic keratoplasty; **healed**, resolution without surgery; **unfavorable course**, prolonged therapy and/or corneal scarring; **variable/indefinite**, outcome uncertain from the current profile and requiring further evaluation.

**Supplementary Figure 1.**

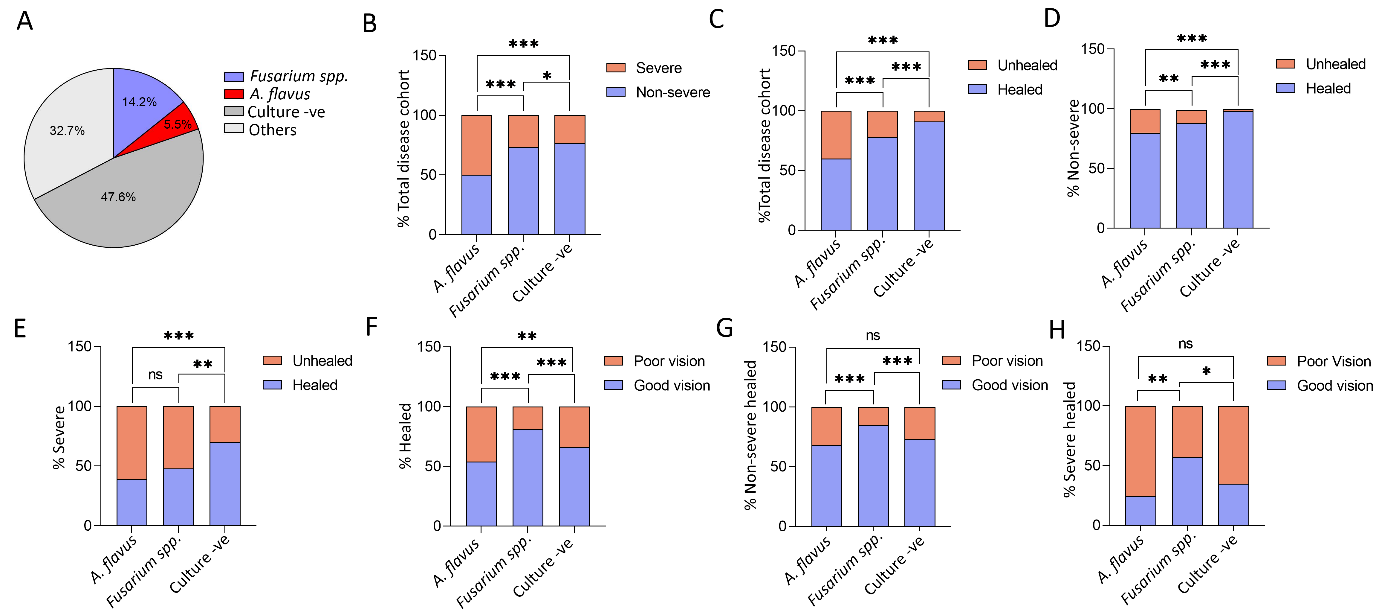

**Supplementary Figure 1. Comparison of clinical presentation and outcomes across etiologies of keratitis.**

Pie chart illustrates the relative proportion of different pathogen reported among keratitis patients during the period of year 2021-2024 at the Cornea Clinic, Aravind Eye Hospital, Madurai **(A).** Stacked bar graphs represent the percentage distribution-Clinical presentation based on ulcer size and depth at the time of initial clinical evaluation and classified into ‘non-severe’ or ‘severe’ **(B)**; Clinical outcome classified based on treatment response as ‘healed’ (infection resolution with corneal healing) or ‘unhealed’ (non-response to medical therapy requiring TPK) **(C-E)**; Visual outcomes among healed cases and classified good vision (complete or improved vision relative to baseline) or poor vision (no improvement or deterioration relative to baseline) **(F-H)**.

**Supplementary Figure 2.**

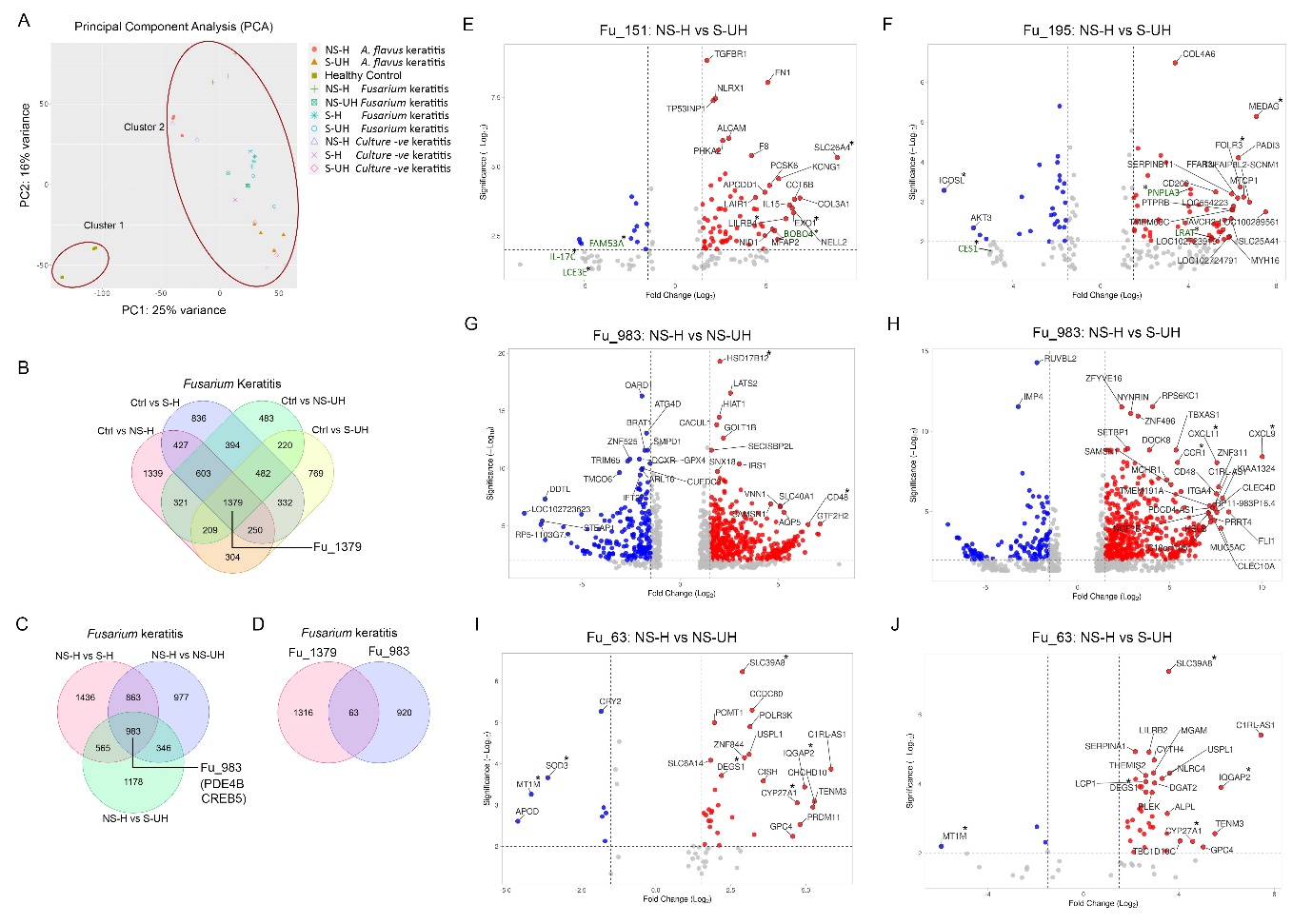

**Supplementary Figure 2. Transcriptomic profiling and identification of host-response gene signatures associated with clinical trajectory in *Fusarium* Keratitis.**

PCA plot of the transcriptome profiles across the different disease groups and healthy controls, illustrating the distinct separation of healthy controls from the diseased cohorts **(A)**. Venn diagram comparing lists of differentially expressed genes (DEGs) in four different sample groups of *Fusarium* keratitis relative to healthy controls; 1379 common DEGs (described as Fu_1379) represent broad disease-associated changes **(B)**. Venn diagram comparing DEGs in three different sample groups of *Fusarium* keratitis relative to the non-severe healed cohort; 983 common DEGs (described as Fu_983) represent changes associated specifically with clinical severity and treatment outcomes **(C)**. Venn diagram comparing the Fu_1379 and Fu_983 gene sets; the common 63 DEGs (described as Fu_63) illustrate the limited overlap between broad disease responses and severity-specific signatures **(D)**. Volcano plots for the 151 and 195 DEGs in the severe unhealed groups relative to the non-severe healed cohort, respectively, from *Fusarium* keratitis analysis showing fold-change in relative expression versus statistical significance; the top 20 predicted DEGs are shown in black font, additional DEGs are indicated in green, and asterisks (**)* indicate candidates selected for validation **(E–F)**. Volcano plots for the 983 DEGs in the non-severe unhealed and severe unhealed groups, respectively, relative to the non-severe healed group, illustrating relative expression and significance; the top 30 predicted DEGs are indicated and asterisks (*) denote candidates for future validation **(G–H)**. Volcano plots for the 63 DEGs in the non-severe unhealed and severe unhealed groups, respectively, relative to the non-severe healed group, illustrating relative expression and significance; the top 20 predicted DEGs are indicated and asterisks (*) denote candidates for future validation **(I–J)**.

**Supplementary Figure 3.**

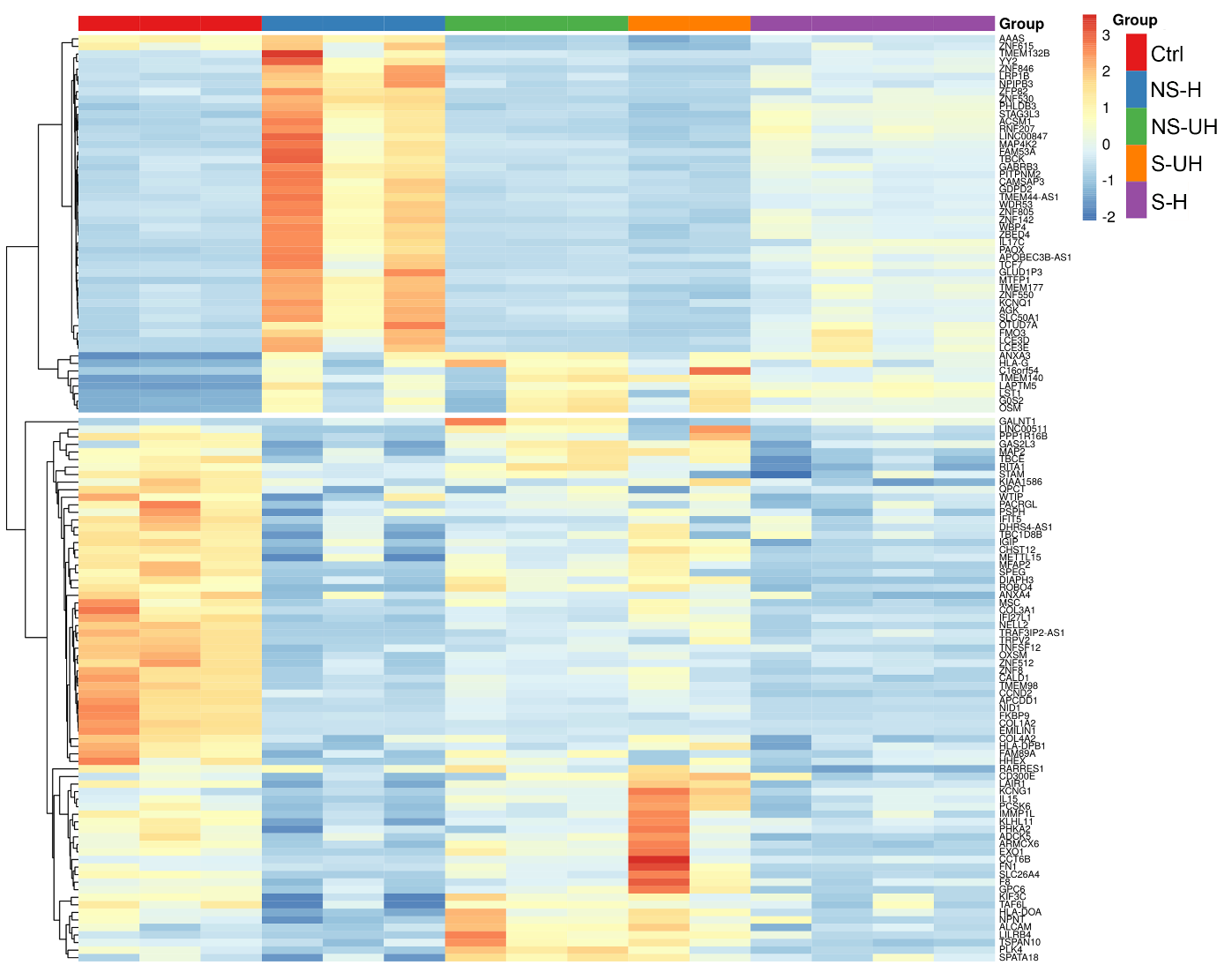

**Supplementary Figure 3. Unsupervised hierarchical clustering of the Fu_151 gene subset in *Fusarium* keratitis showing potential healing-associated signature.**

**Abbreviations:** Ctrl, healthy control (cadaveric cornea); NS-H, non-severe healed; NS-UH, non-severe unhealed; S-H, severe healed; S-UH, severe unhealed.

**Supplementary Figure 4.**

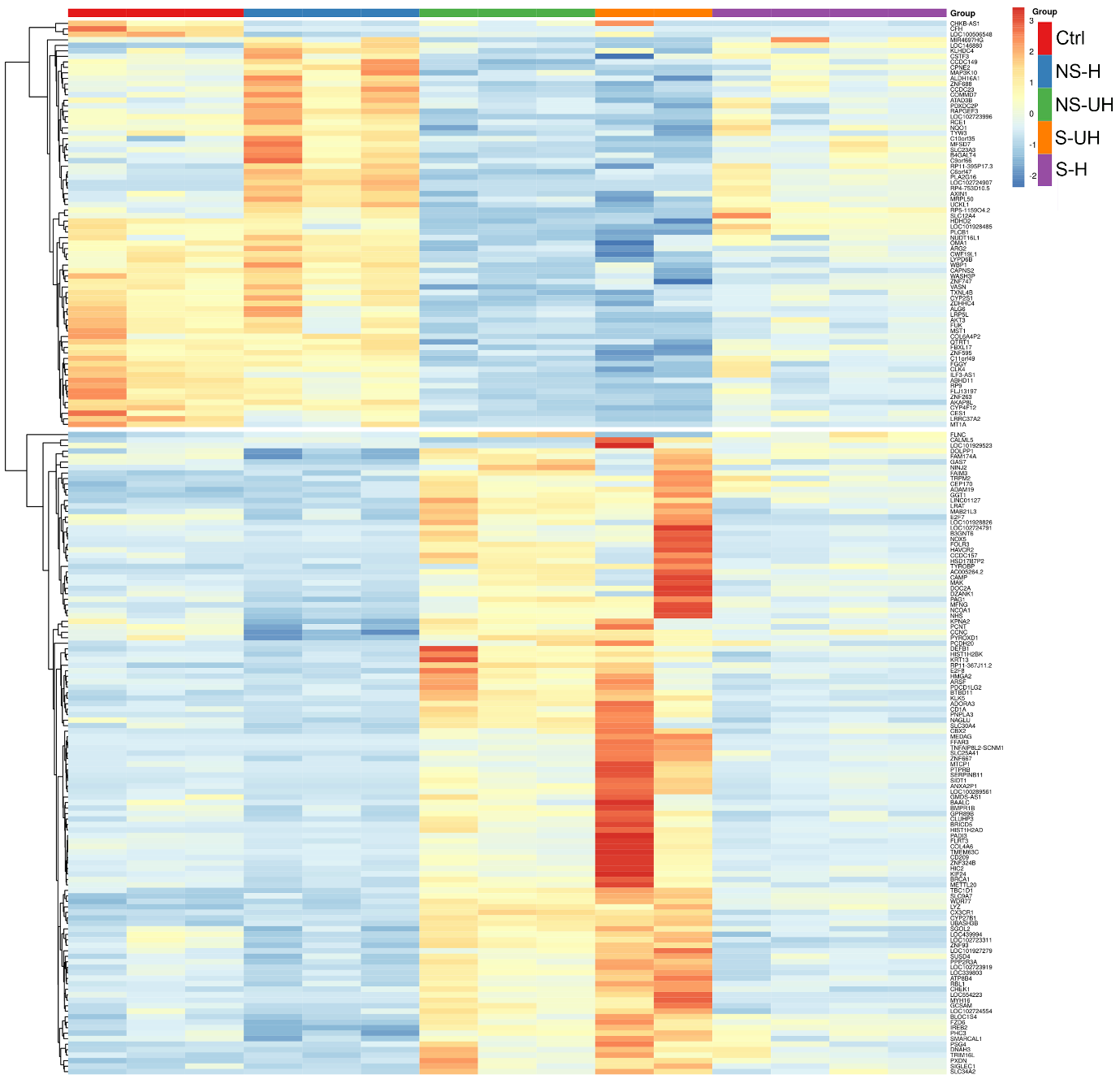

**Supplementary Figure 4. Unsupervised hierarchical clustering of the Fu_195 gene subset in *Fusarium* keratitis showing potential unhealing-associated signature.**

**Abbreviations:** Ctrl, healthy control (cadaveric cornea); NS-H, non-severe healed; NS-UH, non-severe unhealed; S-H, severe healed; S-UH, severe unhealed.

**Supplementary Figure 5.**

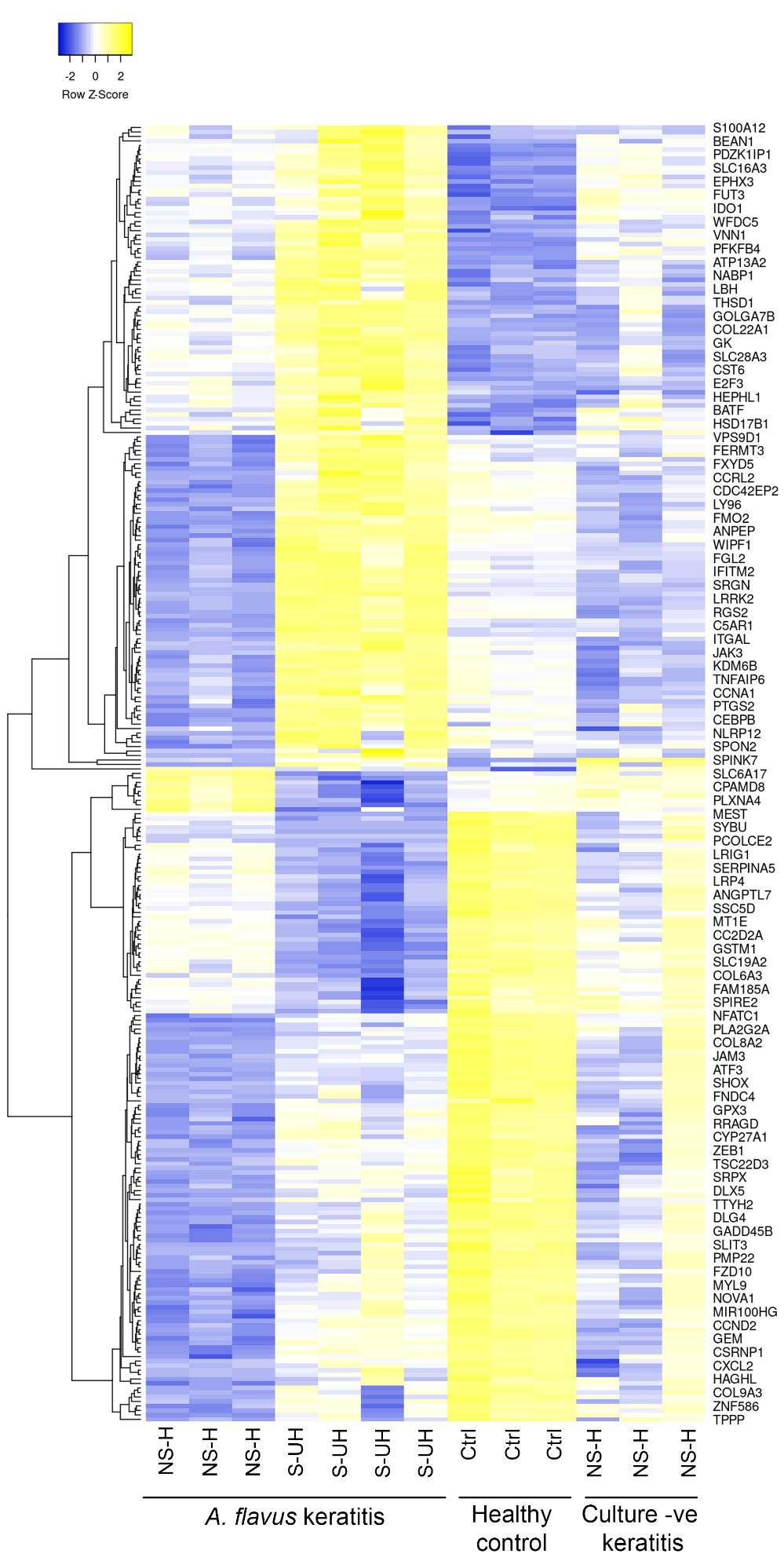

**Supplementary Figure 5. Unsupervised hierarchical clustering of the Afl_287 gene subset in *Aspergillus flavus* showing potential severity/outcome-associated signature.**

**Abbreviations:** Ctrl, healthy control (cadaveric cornea); NS-H, non-severe healed (*A. flavus* keratitis & Culture negative) keratitis); S-UH, severe unhealed.

**Supplementary Figure 6.**

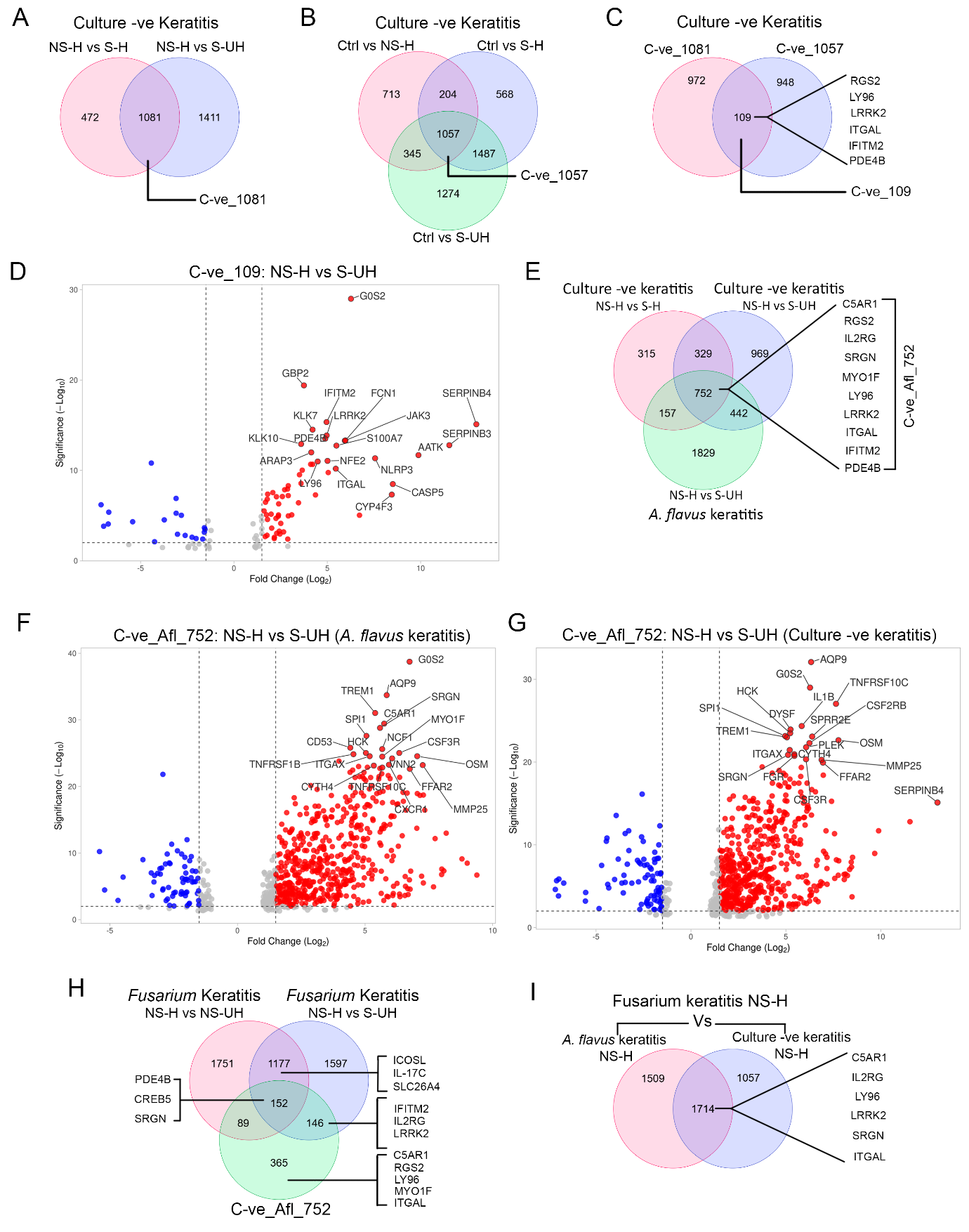

**Supplementary Figure 6. Comparative transcriptomic analysis and identification of shared host-response gene signatures across keratitis etiologies.**

Venn diagram comparing differentially expressed genes (DEGs) in severe healed and severe unhealed groups of culture-negative keratitis relative to the non-severe healed cohort; 1081 common DEGs (described as C-ve_1081) represent signatures associated specifically with clinical severity and treatment outcomes **(A)**. Venn diagram comparing DEGs across all three culture-negative keratitis groups relative to healthy controls; 1057 common DEGs (indicated as C-ve_1057) represent broad disease-associated changes **(B)**. Venn diagram illustrating the overlap between C-ve_1081 and C-ve_1057; the 109 overlapping DEGs (indicated as C-ve_109) contain six candidate severity/outcome-associated genes previously identified in the *A. flavus* keratitis analysis **(C)**. Volcano plot for the C-ve_109 DEGs in the severe unhealed group relative to the non-severe healed group, illustrating the top 20 predicted DEGs, which include five of the six candidate severity signatures from the *A. flavus* keratitis analysis **(D)**. Venn diagram showing the overlap in DEGs between culture-negative and *A. flavus* keratitis samples relative to their respective non-severe healed controls; the 752 overlapping DEGs (indicated as C-ve_Afl_752) contain all candidate severity/outcome-associated genes from the *A. flavus* analysis except *CCRL2* **(E)**. Volcano plots of the C-ve_Afl_752 genes in the severe unhealed group relative to the non-severe healed group for *A. flavus* and culture-negative keratitis, respectively; 13 genes among the top 20 predicted DEGs were shared between etiologies and are indicated by asterisks (*) **(F–G)**. Venn diagram showing the overlap between DEGs in unhealed *Fusarium* keratitis relative to non-severe healed and the C-ve_Afl_752 set, highlighting etiology-specific expression profiles; *PDE4B, CREB5,* and *SRGN* were shared across DEG lists across all keratitis types, while *ICOSL, SLC26A4,* and *IL17C* were differentially expressed in unhealed *Fusarium* groups relative to the non-severe healed; DEGs including *IFITM2, IL2RG,* and *LRRK2* were shared across all severe unhealed etiologies, whereas *C5AR1, RGS2, LY96, MYO1F,* and *ITGAL* were shared by *A. flavus* and culture-negative keratitis but not *Fusarium* keratitis when compared with their non-severe healed counterparts **(H)**. Venn diagram comparing DEGs in the non-severe healed groups of *A. flavus* and culture-negative keratitis relative to the non-severe healed group of *Fusarium* keratitis; 1714 common DEGs distinguish these profiles from *Fusarium* non-severe healed and include six of the selected severity/outcome-associated candidate signatures from the *A. flavus* analysis **(I)**.

**Supplementary Figure 7.**

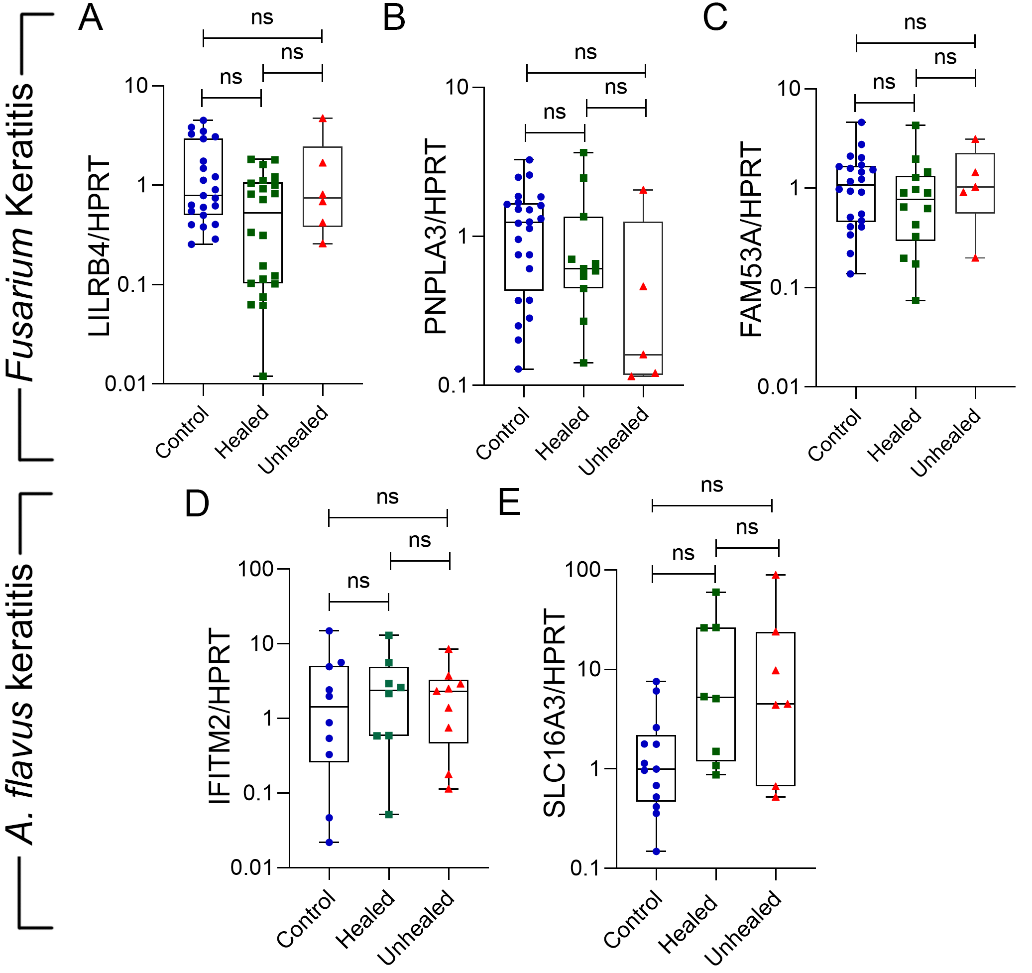

**Supplementary Figure 7.** **Validation of gene expression in *Fusarium* and *A. flavus* keratitis patient samples.**

mRNA expression levels of target genes relative to housekeeping gene HPRT in corneal samples from patients with Fusarium **(A-C)** and A. flavus **(D-E)** keratitis, fold changes calculated over healthy cadaveric controls displayed as box plots showing distribution across control, healed, and unhealed groups. Individual data points are shown, and statistical significance was determined using the nonparametric Kruskal-Wallis test with Dunn multiple comparison: ns, nonsignificant.

**Supplementary Figure 8.**

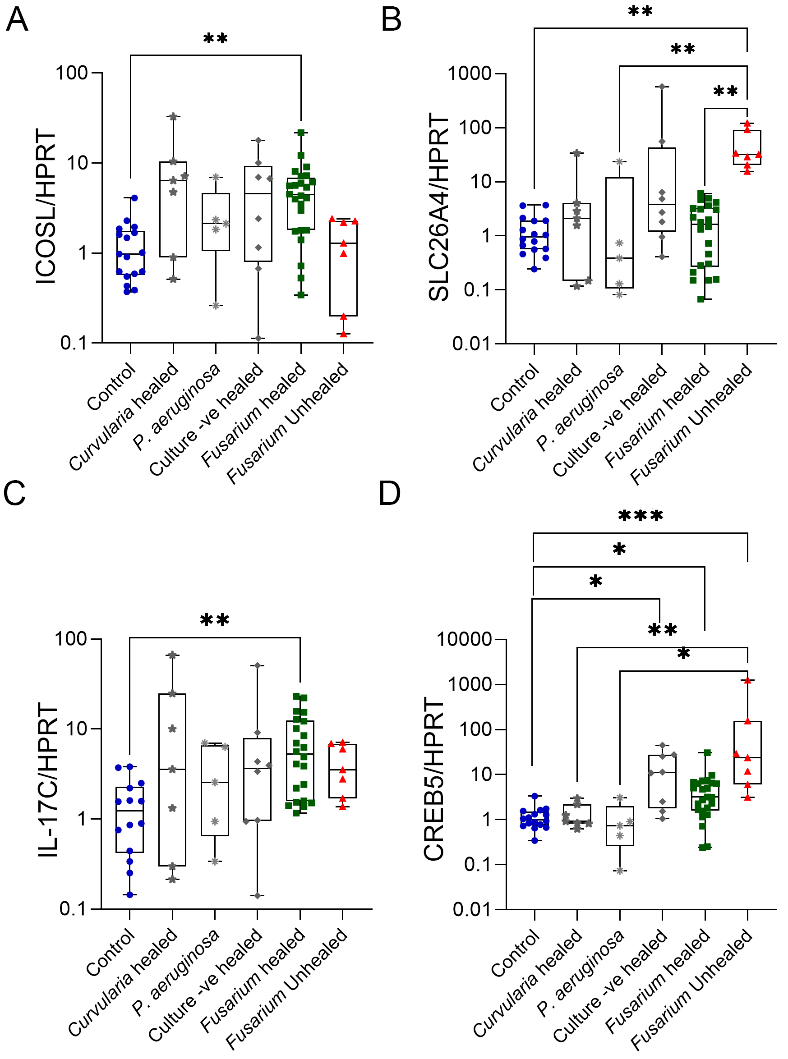

**Supplementary Figure 8. Comparison of outcome-associated gene expression signatures of *Fusarium* keratitis with healed groups of other etiologies.**

mRNA expression levels of target genes relative to housekeeping gene HPRT in corneal samples from patients with *Fusarium* and other keratitis types **(A-D)**, fold changes calculated over healthy controls displayed as box plots showing distribution across control, healed (*Curvularia* spp., *P. aeruginosa*, and *Fusarium* spp.), and unhealed *Fusarium* groups. Individual data points are shown, and statistical significance was determined using the nonparametric Kruskal-Wallis test with Dunn multiple comparison: *P < .05; **P < .01; ***P < .001; ns, nonsignificant.

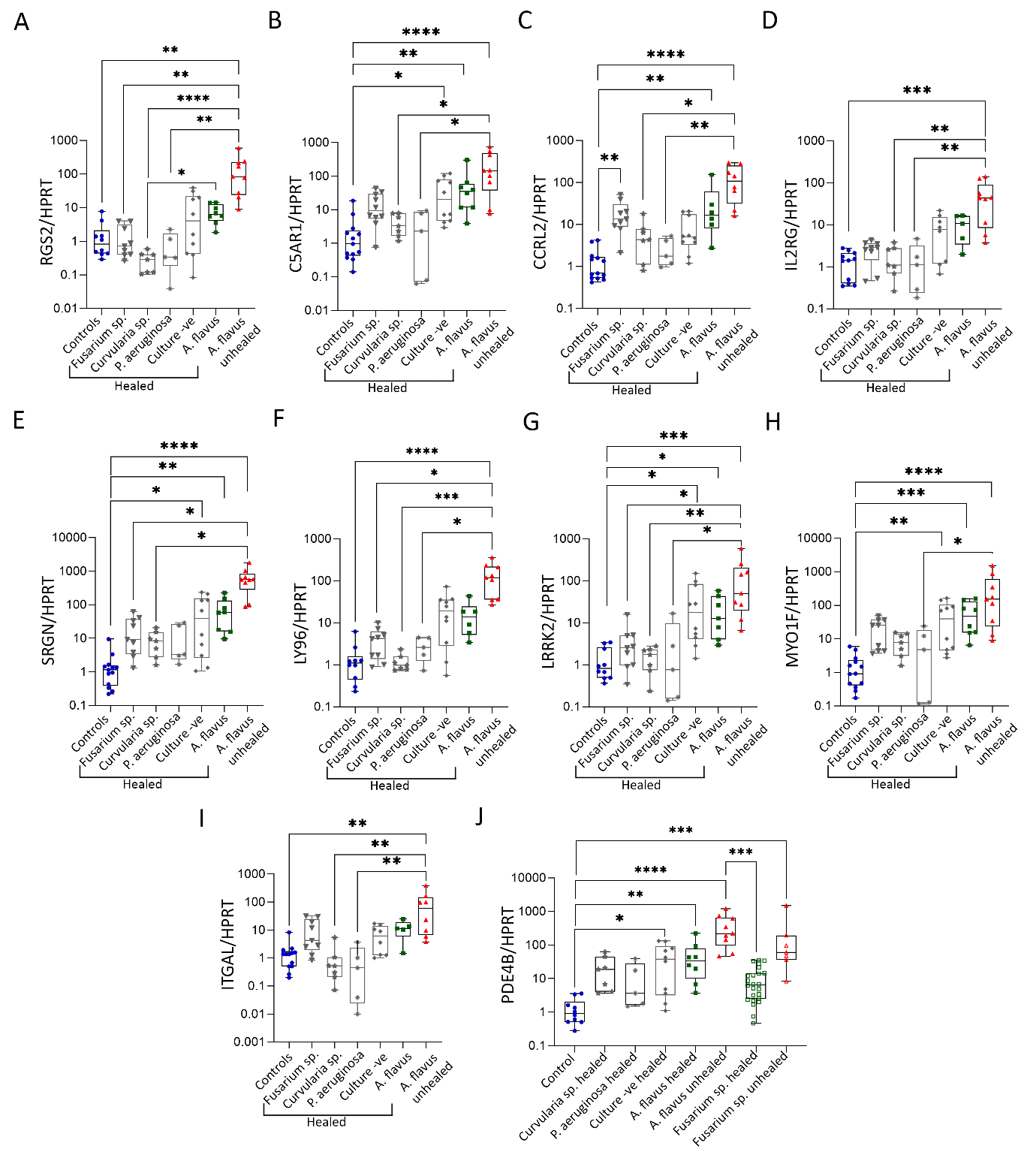
**Supplementary Figure 9.**

**Supplementary Figure 9. Comparison of severity/outcome-associated gene expression signatures of *A.flavus* keratitis with healed groups of other etiologies.**

mRNA expression levels of target genes relative to housekeeping gene *HPRT* in corneal samples from patients with various microbial keratitis types (**A-J**), fold changes calculated over healthy controls displayed as box plots showing distribution across control, healed (*Fusarium* sp., *Curvularia* sp., *P. aeruginosa*, and *A. flavus*), and unhealed (*A. flavus* and *Fusarium* spp.) groups. Individual data points are shown, and statistical significance was determined using the nonparametric Kruskal-Wallis test with Dunn multiple comparison: *P < .05; **P < .01; ***P < .001; ****P < .0001; ns, nonsignificant.

**Supplementary Figure 10.**

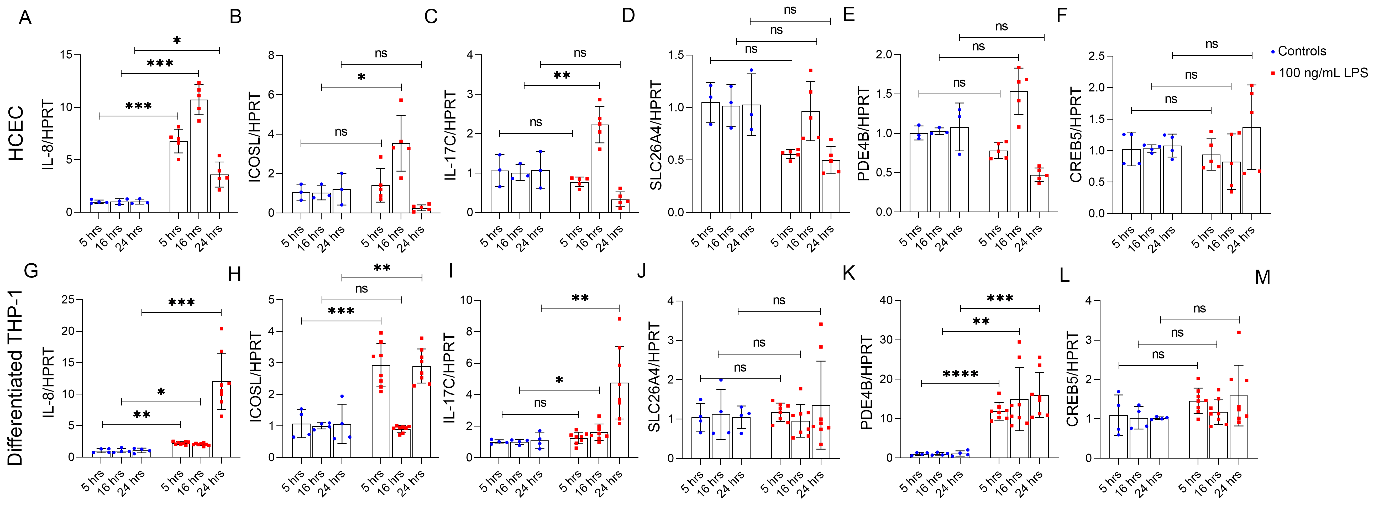

**Supplementary Figure 10. Effect of LPS stimulation on the outcome-associated candidate genes of *Fusarium* keratitis in cell models**.

mRNA expression levels of target genes relative to housekeeping gene HPRT in human corneal epithelial cells (HCECs; **A-H**) and differentiated THP-1 cells **(I-R)** stimulated with 100 ng/mL lipopolysaccharide (LPS) for 5, 16, and 24 hours, fold changes calculated over uninfected controls displayed as bar graphs. Individual data points are shown, and statistical significance was determined using the nonparametric Mann-Whitney U test comparing controls and stimulated groups at each time point: *P < .05; **P < .01; ***P < .001; ****P < .0001; ns, nonsignificant.

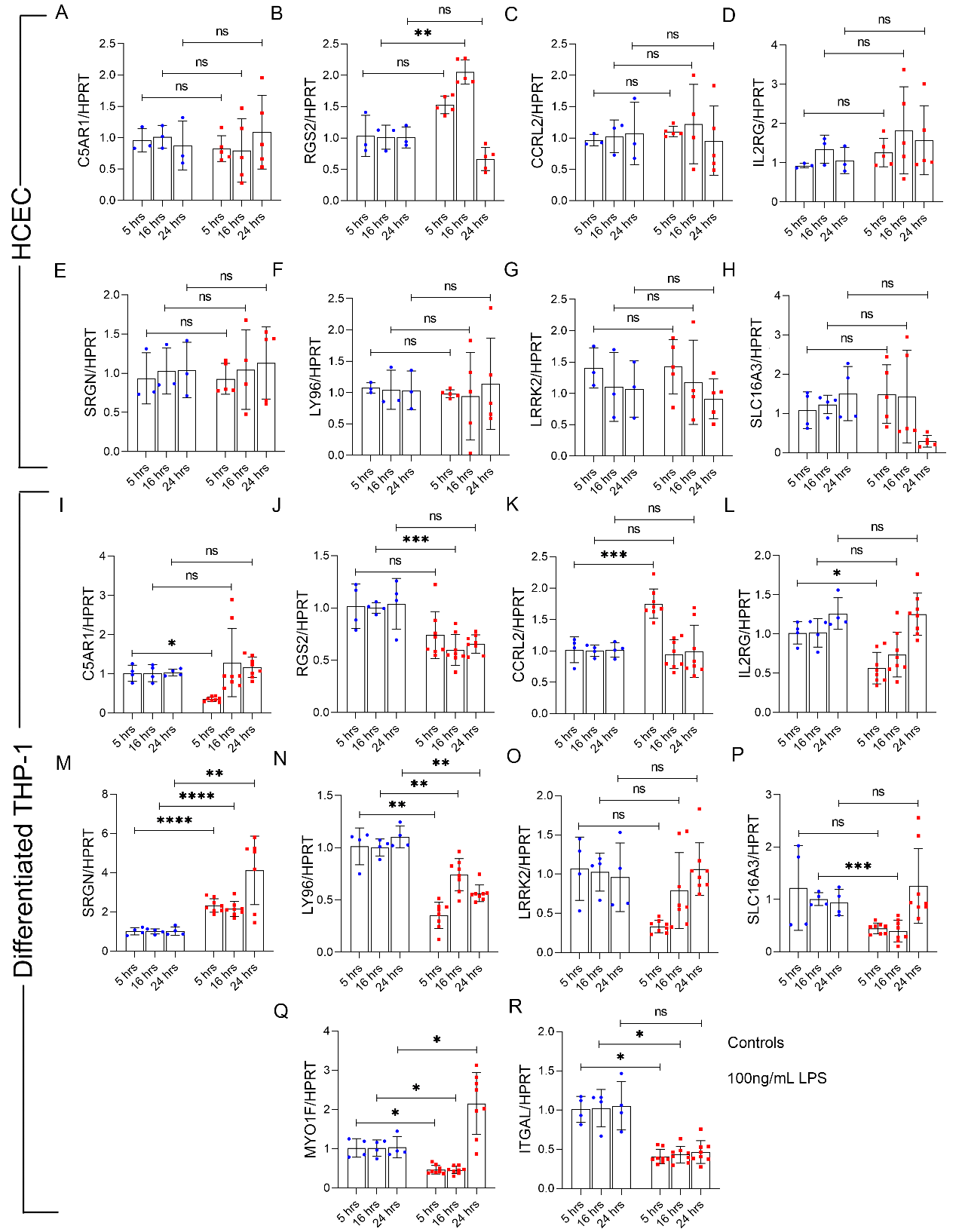
**Supplementary Figure 11.**

**Supplementary Figure 11. Effect of LPS stimulation on the severity/outcome-associated candidate genes of *A.flavus* keratitis in cell models**.

mRNA expression levels of target genes relative to housekeeping gene *HPRT* in human corneal epithelial cells (HCECs; **A-H**) and differentiated THP-1 cells **(I-R)** stimulated with 100 ng/mL lipopolysaccharide (LPS) for 5, 16, and 24 hours, fold changes calculated over uninfected controls displayed as bar graphs. Individual data points are shown, and statistical significance was determined using the nonparametric Mann-Whitney U test comparing controls and stimulated groups at each time point: *P < .05; **P < .01; ***P < .001; ****P < .0001; ns, nonsignificant.

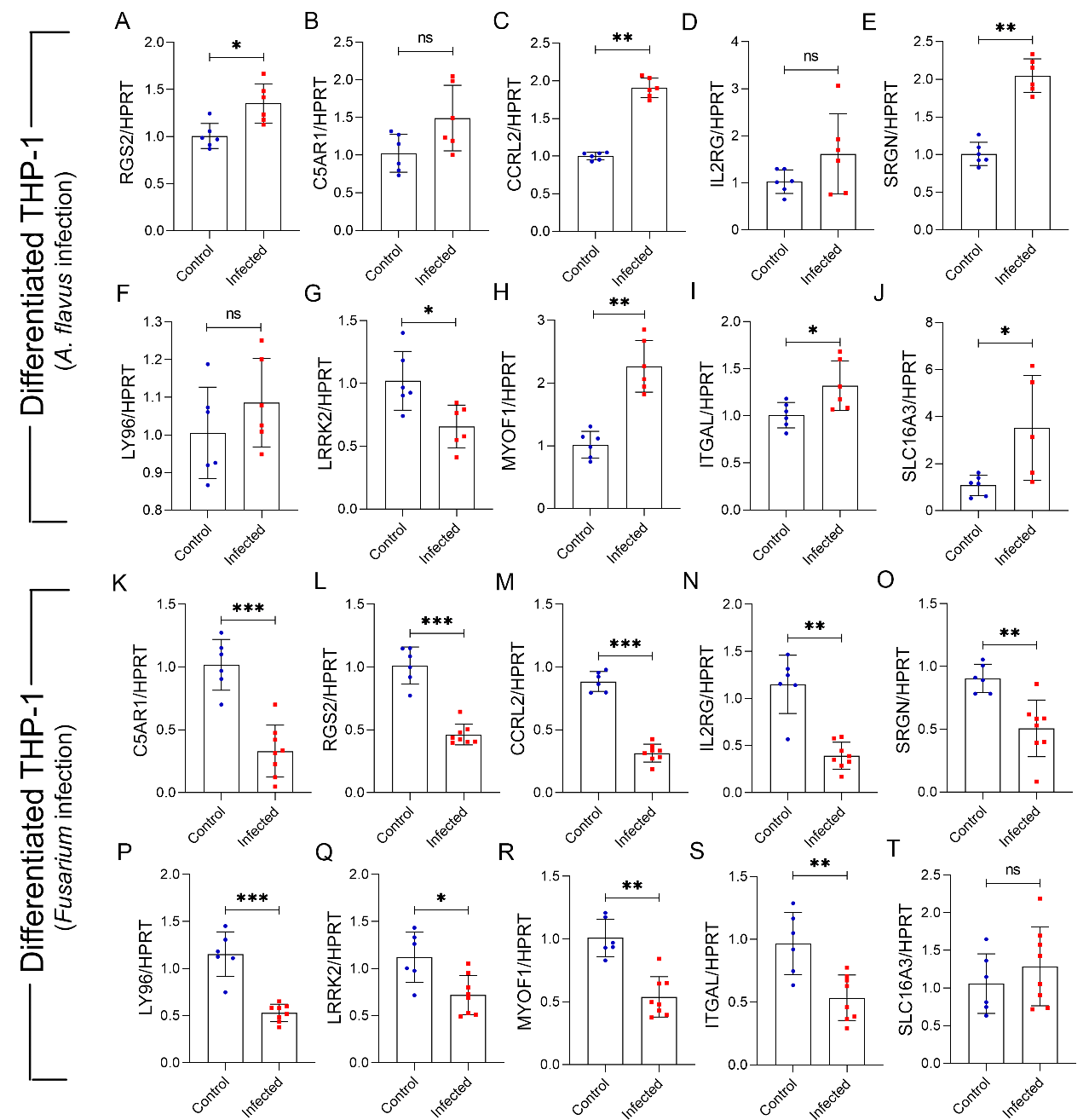
**Supplementary Figure 12.**

**Supplementary Figure 12. Effect of fungal spore co-culture on the severity/outcome-associated candidate genes of *A.flavus* keratitis in cell models.**

mRNA expression levels of target genes relative to housekeeping gene *HPRT* in differentiated THP-1 cells infected with spores of *A. flavus* (**A-J**) and *Fusarium* sp. (**K-T**), fold changes calculated over uninfected controls displayed as bar graphs. Individual data points are shown, and statistical significance was determined using the nonparametric Mann-Whitney U test: *P < .05; **P < .01; ***P < .001; ns, nonsignificant.

**Supplementary Figure 13.**

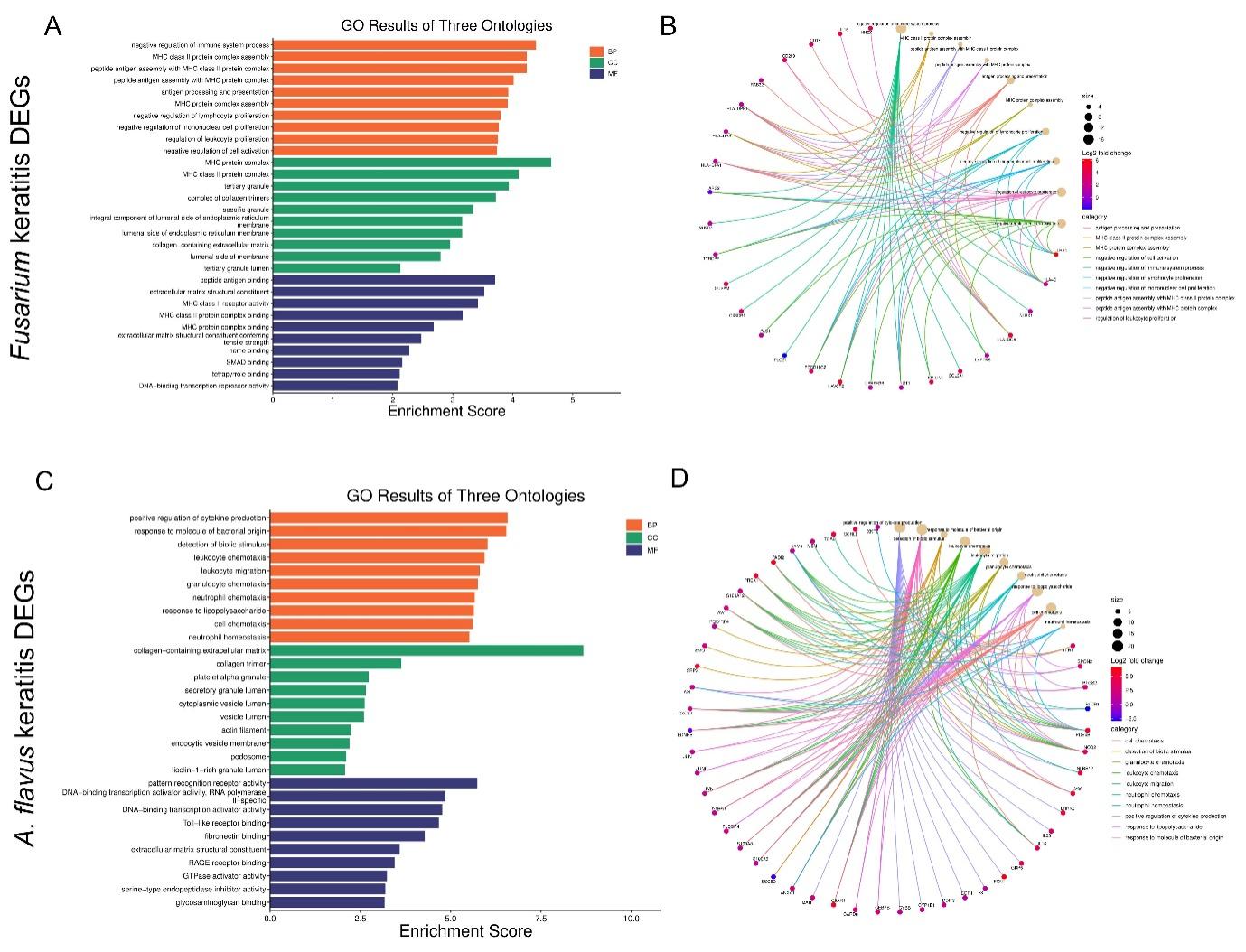

**Supplementary Figure 13. Gene Ontology (GO) enrichment analysis of DEGs from *Fusarium* and *A. flavus keratitis* RNA-seq datasets.**

*Fusarium keratitis* DEGs (Fu_151 + Fu_195) **(A,B)** ; *A. flavus keratitis* Differentially Expressed Genes (DEGs) (Afl_287) **(C,D)**. Bar plots showing enriched GO terms across three ontologies: biological process (BP), cellular component (CC), and molecular function (MF). Bars are ordered by enrichment score, with bar length indicating the magnitude of enrichment. Colors denote GO ontology categories (BP, CC, MF as indicated in the legend) **(A,C).** Gene–concept network plots (cnetplots) illustrating the relationships between enriched GO terms and associated DEGs. Larger nodes represent GO terms, while smaller nodes represent individual genes. Edges indicate gene membership within each GO category. Gene node color reflects log2 fold change, with the color scale indicating relative up- and downregulation **(B,D)**. Enrichment analyses for each dataset were performed independently using sthe same parameters, and only the top enriched GO terms are displayed.
